# Macropinosomes host TLR9 signaling and regulation of inflammatory responses in microglia

**DOI:** 10.1101/2021.02.11.430773

**Authors:** Yu Hung, Neeraj Tuladhar, Zhi-jian Xiao, Samuel J Tong, Jana Vukovic, Lin Luo, Adam A Wall, Jennifer L Stow

## Abstract

To support their innate immune and scavenging functions in the brain, microglia are equipped with Toll-like receptors (TLRs), including the intracellular receptor TLR9, which is activated by microbial CpG-rich DNA. Macropinocytosis is an abundant and inducible pathway in microglia for fluid-phase uptake and ingestion of microbes and cell debris. TLR9 signaling has been ascribed to endolysosomes, particularly lysosomes, which it accesses through direct transport or via internalization from the surface. Here, TLR9 and exogenous CpG-DNA are localized during uptake into fluid-filled macropinosomes, upon upregulated macropinocytosis, where acidic and proteolytic environments support MyD88-induced signaling. Macropinosomes represent an abundant pathway for endolysosomal traffic of TLR9 but are also a much more exposed site for nucleic acid activation of the receptor with a risk of excessive inflammation. To constrain TLR9 inflammation, macropinosomes also house the TLR9 co-receptor LRP1 and regulators Rab8a and PI3Kγ which augment Akt signaling and favor anti-inflammatory cytokine production. Macropinosomes and their inflammatory regulators are therefore important components of TLR9 pathways in microglia that are poised for surveillance and protection in the CNS.

## Introduction

As a specialized population of innate immune cells in the central nervous system (CNS), microglia maintain brain homeostasis by supporting neuronal growth and offering surveillance and defensive responses to injury and infection (Wolf, Boddeke et al., 2017). Microglial cells are equipped with pattern recognition receptors, including members of the Toll-like receptor (TLR) family, which detect a wide range of pathogen signatures and signals from cell damage (Block, Zecca et al., 2007, Olson & Miller, 2004). The activation of TLRs triggers innate immune and inflammatory responses, including the secretion of cytokines and chemokines that can mount protective responses to ward off infection (Akira & Takeda, 2004). Microglia are also scavengers, helping to clear the CNS of microorganisms, cell debris, protein aggregates or particles (Davalos, Grutzendler et al., 2005, Nimmerjahn, Kirchhoff et al., 2005) which are normally ingested and degraded, although notable exceptions are the abnormal protein aggregates that infamously accumulate as hallmarks of neurodegenerative disease (Neumann, Kotter et al., 2009). Inflammation triggered inappropriately or in excess, by pathways including TLRs, can contribute to acute and chronic inflammation in the CNS, including in chronic neurodegenerative conditions such as Alzheimer’s and Parkinson’s diseases (Cherry, Olschowka et al., 2014, Perry, Nicoll et al., 2010).

Among members of the TLR family, TLRs 3, 7, 8 and 9 are characterized as intracellular receptors which typically signal from endolysosomal compartments, with a remit to detect and respond to intracellular viral and bacterial pathogens (Lee & Barton, 2014). TLR9 recognizes DNA, particularly unmethylated DNA rich in CpG repeats, characteristic of microbial genomes, but which is also a feature of host mitochondrial DNA (mtDNA). Limiting TLR9 responses to host DNA is critical for avoiding excessive inflammation and autoimmune disease (Barton & Kagan, 2009, Barton, Kagan et al., 2006). The intracellular location of TLR9 and sequestration of its activation and signaling environments have been proposed as a major adaptation designed to limit the scope of TLR9 activation and responses (Barton & Kagan, 2009).

TLR9 accumulates in the endoplasmic reticulum (ER) in dendritic cells (Combes, Camosseto et al., 2017, Latz, Schoenemeyer et al., 2004, Leifer, Kennedy et al., 2004) and in lysosomes (Ahmad-Nejad, Häcker et al., 2002, Häcker, Mischak et al., 1998), the latter of which has been designated as the major compartment for TLR9 activation and signaling. The lysosomal milieu is an acidic and proteolytic environment which can release nucleic acids from microbes and also support the acid-dependent cleavage of the TLR9 ectodomain as a prerequisite for its ligand- and adaptor-mediated signaling (Ewald, Engel et al., 2011, Ewald, Lee et al., 2008, Häcker et al., 1998, Latz, Verma et al., 2007, Park, Brinkmann et al., 2008). However, TLR9 is also trafficked to the cell surface from where it can access endosomal pathways for internalization, with the potential for signaling in earlier endosomal compartments (Lee, Moon et al., 2013). The identity of pathways responsible for TLR9 internalization and the overall spatiotemporal nature of TLR9 and CpG-DNA trafficking, engagement and signaling are yet to fully understood, especially in different cell types. TLR9 is expressed in microglia, and although it has been implicated in neuroinflammation and neurodegenerative disease, relatively little is known about the receptor’s trafficking or signaling in these cells.

Microglia are exposed to high levels of neuronal cell death in brain injury (Willis, MacDonald et al., 2020) and in ageing brains, where cell debris and nucleic acids are actively scavenged by microglia.

Cell debris and microorganisms are routinely scavenged by macropinocytosis (Canton, 2018, Mercer & Helenius, 2009, Stow, Hung et al., 2020). Macropinosomes are large fluid-filled endocytic compartments derived from closure of cell surface ruffles (Kerr & Teasdale, 2009, Swanson, 2008) that mature to direct cargo into endolysosomal and recycling pathways. In macrophages, macropinocytosis is constitutive, and further upregulated by growth factors or TLRs, constituting the major pathway for abundant fluid-phase uptake and membrane turnover in these cells (Steinman, Brodie et al., 1976, Stow et al., 2020). In such cells, with very active macropinocytosis, this route constitutes the most abundant portal for delivering soluble or membrane-attached cargo to lysosomes. Finally, macropinosomes are also signaling hubs, hosting signaling GTPases and kinases through enrichment of lipids including, PI(3,4,5)P_3_ (Buckley & King, 2017, Yoshida, Hoppe et al., 2009). As a basis for our current investigation, macropinosomes have previously been shown as sites for signaling from TLR4 in macrophages (Wall, Condon et al., 2019, Zanoni, Ostuni et al., 2011). Based on their abundance of transport and their signaling capacity, macropinosomes have been mooted as possible sites for TLR9 signaling (Canton, Schlam et al., 2016, Wall, Luo et al., 2017) but the evidence to support this notion has not been formally presented.

Finally, the significance of macropinosomes as a potential site for TLR9 signaling also relates to molecular regulators that operate in macropinosomes to augment PI3K/Akt signaling and serve as key constraints on TLR-induced inflammation. The class 1B PI3K, PI3Kγ is recruited downstream of multiple TLRs for Akt signaling serving to bias macrophage programming and enhance anti-inflammatory cytokine responses (Kaneda, Messer et al., 2016, Luo, Wall et al., 2014, Schmid, Avraamides et al., 2011). PI3Kγ is recruited as an effector by the activated form of the GTPase Rab8a, which in turn is recruited to TLR signaling pathways by the endocytic receptor LRP1, also a TLR co-receptor (Luo, Wall et al., 2018). Such regulators potentially have key roles in balancing pro- and anti-inflammatory outcomes in disease (Kaneda et al., 2016). The control and influence of inflammatory mediators in microglia is critically important but also complex, as highlighted recently by the unexpected role of pro-inflammatory mediators such as IL-6, which helps to promote recovery from traumatic brain injury (Willis et al., 2020).

The findings presented herein make the case for macropinocytosis as an active pathway in microglia, serving for the uptake of CpG-rich DNA, as well as for the localization of TLR9. We demonstrate the suitability and efficacy of macropinosomes as requisite signaling locales for TLR9 and we show that the LRP1/Rab8a/PI3Kγ complex offers regulation of macropinosomal signaling and modulates inflammatory outputs in microglia.

## Results

### Microglia upregulate macropinocytosis in response to TLR9 activation

In macrophages, ruffling and macropinocytosis occur constitutively and are known to be further enhanced by TLR activation (Canton et al., 2016, Wall et al., 2017). We examined this response in labeled primary microglia in astroglial co-cultures derived from transgenic mice (Fig 1). Cells were seen to constitutively produce lateral and dorsal F-actin-rich ruffles and the number of dorsal ruffles increased significantly after cells were exposed to the TLR9 ligand unmethylated CpG-DNA (Fig 1A). Live imaging of the primary microglia demonstrates that ruffling can result in the formation, closure and internalization of vacuolar-like macropinosomes that are initially distinctly larger than other endosomes (Fig 1B). Incubation of these cells with 488-dextran as a fluid phase marker, demonstrates macropinocytic uptake and measurements of fluorescence intensity show increased uptake of dextran in CpG-DNA exposed cells (Fig 1C). Dextran uptake was then performed on the microglial-like cell line BV-2, before and after exposure to TLR ligands, to visualize and quantify macropinocytosis using an automated image-based program (Condon, Heddleston et al., 2018). The results confirm that CpG-DNA-induces significant increases in total dextran uptake, as well as increases in macropinosome number per cell and size (Fig 1D, Appendix Fig S1). Activated TLR9 more robustly induces macropinocytosis compared to Poly I:C activation of TLR3, as another intracellular TLR in these cells. Hence, in microglia, macropinocytosis is an active endocytic pathway that is strongly increased by TLR9 activation.

**Figure 1.**
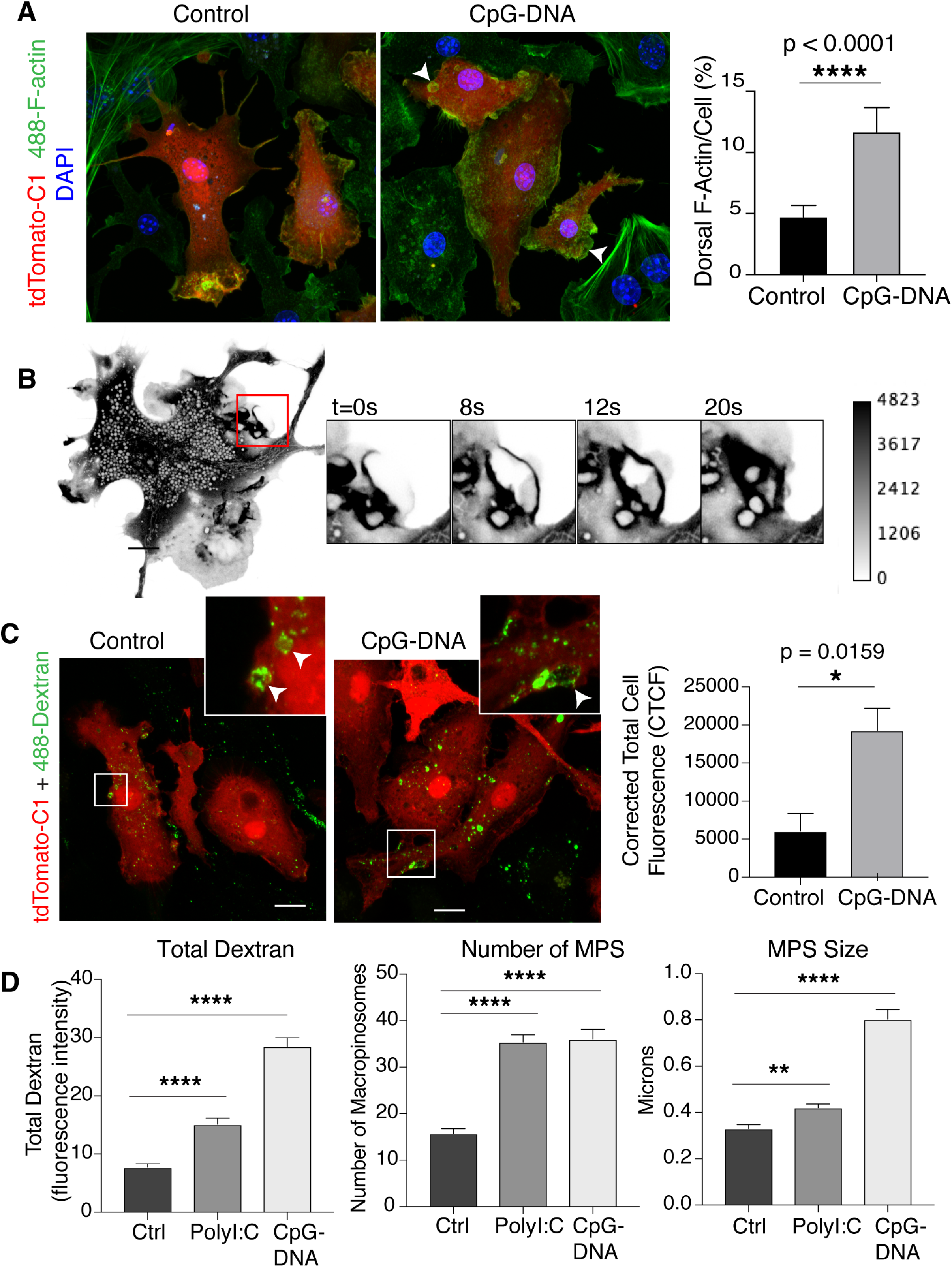
Ruffling and macropinocytosis in microglia. **A**. In primary astroglial cultures, microglia are denoted by expression of cytoplasmic tdTomato-C1 (red) (cultured from CX3CR1creERT2 x iDTR x tdTomato mice); cells are fixed and stained with phalloidin (green) to depict F-actin-rich cell surface projections and ruffles, nuclei are labeled with DAPI (blue). Ruffling was observed and measured in cells treated with CpG-DNA compared to untreated controls. Arrows point to F-actin ruffles. F-actin-staining intensity was measured relative to cell staining in maximum intensity projections. **B**. Confocal live imaging of a representative primary microglial cell expressing cytoplasmic tdTomato-C1 (shown as grey scale) depicts dynamic ruffling and documents the formation and internalization of vacuolar macropinosomes from closure of ruffles. **C**. Primary microglia expressing tdTomato-C1 (red) were incubated with the fluid-phase marker, fluorescent dextran (488-dextran 70K MW, green) for 15 mins, then fixed. The corrected total cell fluorescence (CTCF, which corrects the fluorescence against the background) of dextran per cell was quantified by hand. \**p = 0*.*0159* (unpaired, two-way T-test). **D**. Quantification of dextran staining using an automated script to determine total dextran internalized, number of macropinosomes and mean macropinosome size per cell in BV-2 microglia activated with endosomal TLR ligands: Poly I:C and CpG-DNA. Macropinosome abbreviated to MPS. Cells were incubated with 488-dextran (green) for 1 hour, then fixed and stained with wheat germ agglutinin (red) to define cell outlines. *\*\*\*\*p<0*.*0001* across all graphs (ordinary one-way ANOVA with Dunnett’s multiple comparisons test, significance between control and Poly I:C/CpG-DNA displayed on graph). Data information: Microscopy images are representative of total populations. Graphs are displayed as mean ± SD. For statistical analysis, pooled measurements from 3 technical replicates of n=3 independent experiments were used. For experiments involving primary cells, cells were taken from n=3 mice and technical measurements from individual cells were pooled. All scale bars represent 10μm.

### TLR9 and CpG-DNA are internalized by macropinocytosis

The role of macropinocytosis in TLR9 localization and CpG-DNA internalization remains largely undisclosed, since earlier studies focused on endolysosomal compartments and receptor-mediated uptake of CpG-DNA by cell surface scavenger receptors in clathrin-coated vesicles (Jozefowski, Sulahian et al., 2006, Latz et al., 2004). We first examined the relationship of TLR9 with macropinosomes by localization of recombinant TLR9 in BV-2 cells, noting that TLRs are notoriously difficult to detect by immunostaining or by expression of tagged receptors. Nevertheless, BV-2 cells were transfected with mCherry-TLR9 with or without co-transfection of GFP-Rab8a which we have previously characterized as a prominent marker of early macropinosomes and macropinosome-derived tubules in macrophages (Wall et al., 2019, Wall et al., 2017). Live cell imaging of mCherry-TLR9 showed stable staining of puncta and endolysosomes in the cytoplasm. In this imaging, no accumulation of TLR9 was detected on the cell surface nor in the endoplasmic reticulum, which is a large repository in dendritic cells but not necessarily in B cells (Chaturvedi, Dorward et al., 2008, Leifer et al., 2004) (Fig 2A). In co-transfected cells, large, mobile macropinosomes at the cell periphery were co-labeled for circumferential GFP-Rab8a, overlapping with punctate TLR9 labeling. TLR9 did not appear on GFP-Rab8a-labeled tubules emanating from ruffles or macropinosomes and GFP-Rab8a was not present on other intracellular TLR9-labeled vesicles (Fig 2B). Thus, we provide evidence for mCherry-TLR9 localization and internalization into macropinosomes.

**Figure 2.**
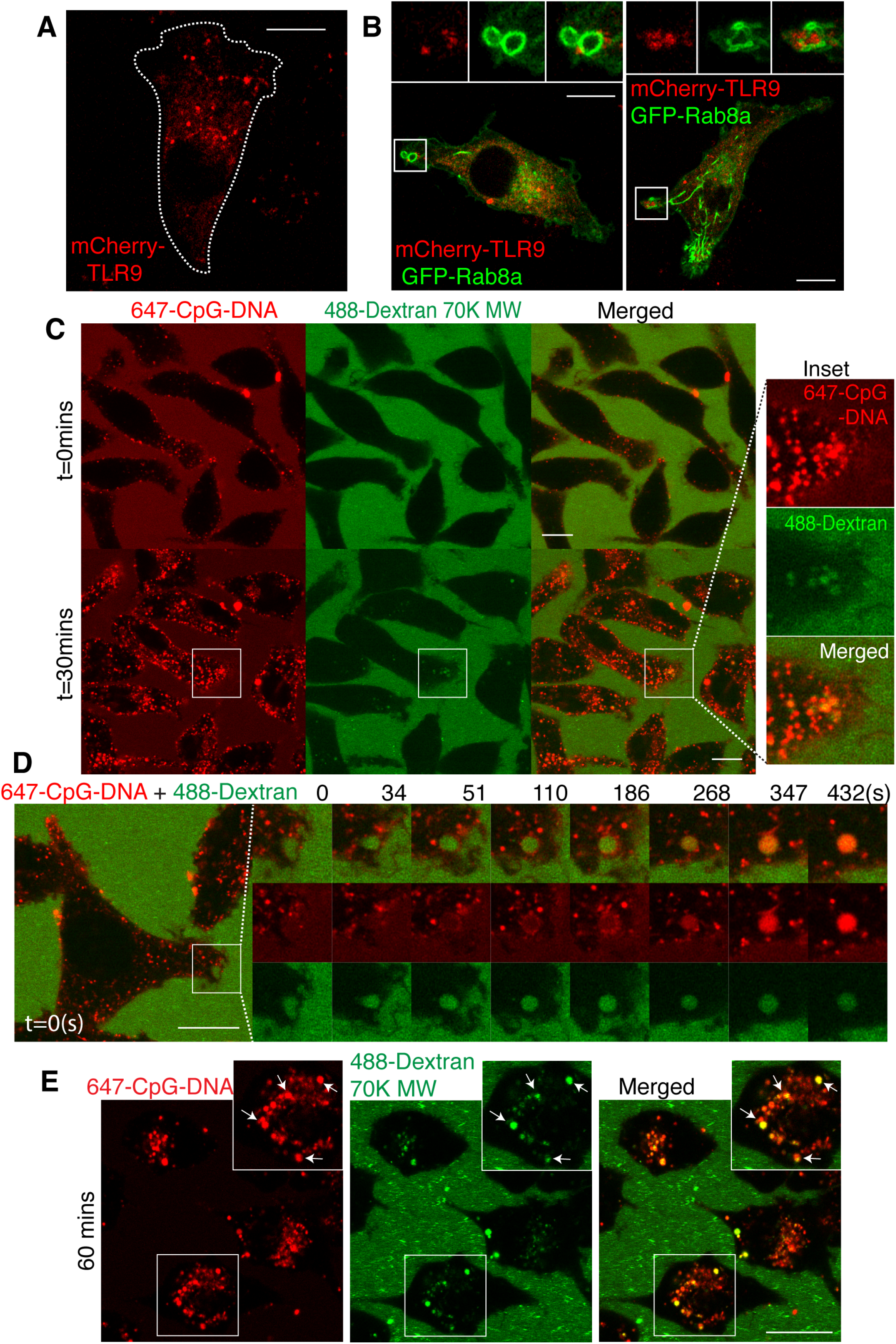
CpG-DNA DNA and TLR9 are internalized in macropinosomes. **A**. Live cell imaging of BV-2 microglia transiently transfected with mCherry-TLR9 (red). Cell outline marked by white dotted line. **B**. Live cell confocal imaging of BV-2 cells transiently transfected with mCherry-TLR9 (red) and GFP-Rab8a (green). GFP-Rab8a prominently outlines early macropinosomes and tubules emanating from them. mCherry-TLR9 is found as punctate labeling, some of which coincides with Rab8a on macropinosomes. **C**. BV-2 cells (appearing in negative relief) are shown incubated with 647-CpG-DNA (red) and 488-dextran 70K MW (green) fluorescing in the medium. Single frames from live confocal imaging recorded immediately upon addition of the labels and after 30 mins when both dextran and CpG-DNA have been internalized. **D**. Movie sequence tracking both fluid-phase 488-dextran 647-CpG-DNA in the lumen of an early macropinosomes. 647-CpG-DNA can be seen in puncta around the macropinosome membrane which pinch off, but also as soluble cargo in the macropinosomes lumen. **E**. 647-CpG-DNA (red) and 488-Dextran 70K MW (green) in live BV-2 microglia after 60 mins uptake. Data information: All scale bars represent 10μm.

In earlier reports, labeled, extracellular CpG-DNA has been shown to be captured by surface scavenger receptors and non-selectively internalized by several cell types; here we sought to more specifically define its uptake. BV-2 cells were incubated in medium containing fluorescent CpG-DNA (647-CpG-DNA) together with 488-dextran. Live uptake of these markers was optimally visualized using cells bathed continuously in the fluorescent medium, wherein the cells themselves are shown in relief (Fig 2C). Immediately after addition, some 647-CpG-DNA is found at the cell surface and is taken up into peripheral macropinosomes also containing 488-dextran. After 30 mins there is a marked accumulation of CpG-DNA in endosomal compartments, some of which are co-labeled with dextran, overall representing intake of CpG-DNA into the endosomal system via macropinosome maturation and possibly by additional endocytic pathways (Fig 2C).

Further inspection of very early macropinosomes reveals that 647-CpG-DNA is captured along with dextran as fluid phase cargo where both are present in the macropinosome lumen, initially at intensities equating to the extracellular medium, consistent with the non-selective gulping that is characteristic of macropinocytosis (Fig 2D). In addition, there is membrane-bound 647-CpG-DNA internalized into early macropinosomes lining the inner surface of these vacuoles where it appears as concentrated puncta, which then bud off (Fig 2D). The maximum intensity of lumenal dextran (at 110s in Fig 2D) precedes a notable concentration of 647-CpG-DNA (from 347s) along with faint tubulation, which could represent concentration and sorting of the CpG-DNA when macropinosome shrinkage is occurring. 647-CpG-DNA was also examined in GFP-Rab8a expressing cells. Both fluid-phase and membrane-bound punctate CpG-DNA were found on Rab8a positive macropinosomes and temporally, the CpG-DNA accumulated beyond the transient residence of GFP-Rab8a (Appendix Fig S2).

At later times (60 mins) both 647-CpG-DNA and 488-dextran are found together in large perinuclear compartments, as a consequence of abundant transport through macropinosomal pathways into endolysosomal compartments (Fig 2E). Staining with Lamp1 confirms the identity of these 647-CpG-DNA compartments as lysosomes (Appendix Fig S3). Taken together, the imaging portrays internalization of both fluid-phase and membrane-bound CpG-DNA into early macropinosomes. From this compartment CpG-DNA accesses later Lamp1-positive endolysosomal compartments which can occur through either bulk maturation of macropinosomes or possibly via receptor-mediated pathways emanating from macropinosomes. Based on the upregulation of macropinocytosis upon TLR9 activation and the high intrinsic capacity of this pathway for membrane turnover, as well as for fluid and debris intake, our evidence points to macropinosomes as a major, initial portal for CpG-DNA uptake and a location for TLR9.

### Inhibition of macropinocytosis blocks CpG-DNA-DNA internalization and TLR9 signaling

The contribution of macropinocytosis to CpG-DNA uptake and to TLR9 signaling were tested in cells treated with the commonly-used macropinocytosis inhibitor, 5-(N-Ethyl-N-isopropyl)amiloride (EIPA) (Koivusalo, Welch et al., 2010). In classical fashion, EIPA treatment dramatically reduced dextran uptake in BV-2 cells (Appendix Fig S4). Under these conditions, 488-dextran and 647-CpG-DNA were visualized and quantified at 15 and 60 mins and inhibition of macropinocytosis dramatically reduced the staining of CpG-DNA and dextran in macropinosomes and endolysosomes at early and later times. Fluorescence intensities per cell were used to measure uptake showing that EIPA nearly ablated intracellular dextran at both times and it reduced intracellular CpG-DNA by over 50% at early and late times (Fig 3). This confirms that macropinocytosis is a predominant route for CpG-DNA uptake into microglial cells and a main route for accumulating CpG-DNA in endolysosomes.

**Figure 3.**
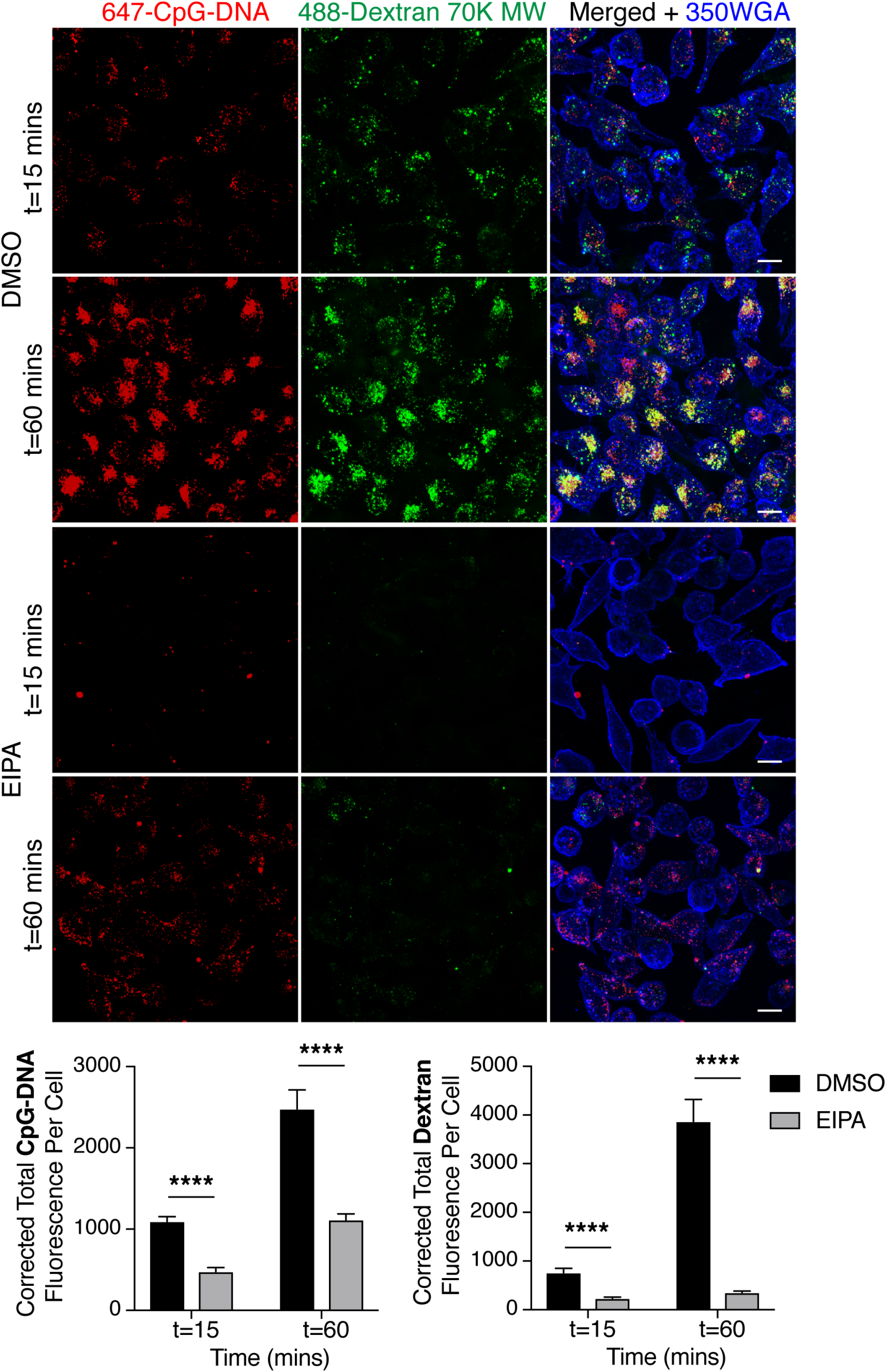
Inhibition of macropinocytosis by EIPA, blocks uptake of CpG-DNA. Max intensity projection of 647-CpG-DNA (red) and 488-Dextran (green) uptake in fixed BV-2 microglia. Total CpG-DNA and dextran fluorescence per cell was captured and quantified at 15 and 60 mins. \*\*\*\**p<0*.*0001* (unpaired, two-tailed Mann-Whitney test). n=5 independent experiments. All scale bars represent 10μm. All graphs are presented as mean ± SD.

The presence of both TLR9 and CpG-DNA in macropinosomes creates a potential niche for receptor activation and signaling. TLR9 signaling was tested in BV-2 cells before and after exposure to CpG-DNA. TLR9 signals from endosomal membranes through the adaptor MyD88 which culminates in proinflammatory cytokine production via MAPK pathways and signaling through the transcription factor NF-kB (Akira & Takeda, 2004). CpG-DNA-treated BV-2 cells recorded phosphorylation of p65, an NF-kB subunit and some activation of ERK1/2. CpG-DNA also induces phosphorylation of Akt; this was first detected between 5 and 15 mins, at a time when our staining indicates that TLR9 and CpG-DNA are in early macropinosomes (Fig 4A). Both MyD88-dependent NF-kB and Akt signaling were markedly attenuated in cells treated with EIPA, confirmed by quantification of the phospho-Akt and phospho-p65 responses.

**Figure 4.**
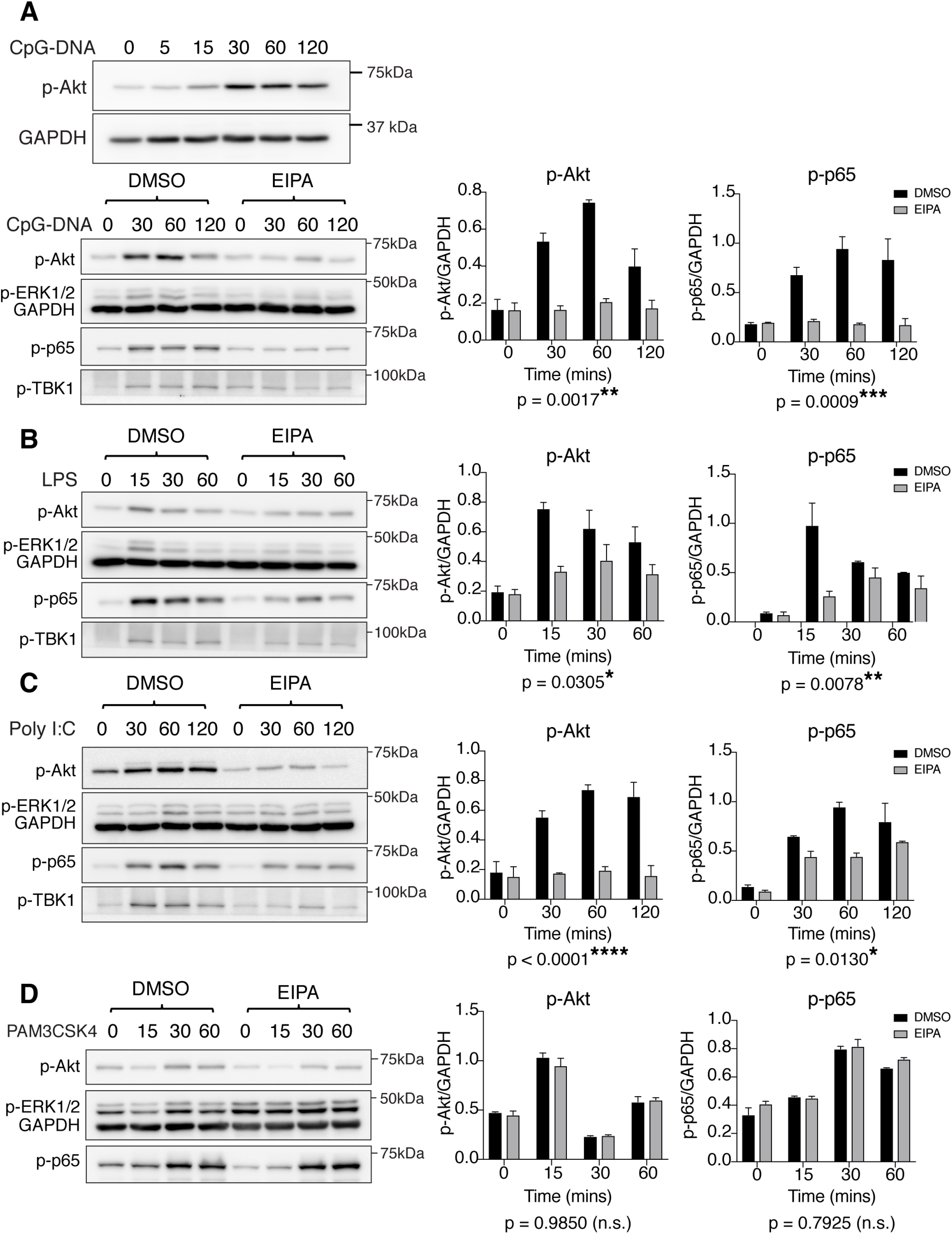
Macropinocytosis is required for TLR9 signaling. **A**. BV-2 microglia activated with different TLR ligands: CpG-DNA **(A)**, LPS **(B)**, Poly I:C **(C)** and PAM3CSK4 **(D)** to examine the phosphorylation of signaling kinases, phospho (p)-Akt, phospho (p)-ERK, phospho (p)-p65 and phospho (p)-TBK1 by immunoblotting on cell extracts. **A)** also includes a short time course for activation, blotting for phospho (p)-Akt. Cells are incubated with DMSO control or 75μM EIPA for 1 hour prior to activation. Representative immunoblots are shown, GAPDH is a loading control. Graphs are displayed as mean ± SD. Two-way ANOVA was used to assess significance, significance for interaction is marked underneath graph. n=3 independent experiments for all panels.

For comparison, signaling generated by activation of other TLRs was examined. LPS activation of TLR4 resulted in Akt and p65 responses that were only partially blocked by EIPA, in keeping with well-known TLR4 signaling generated at early times from both the cell surface and macropinosomes (Kagan, Su et al., 2008, Luo et al., 2014) (Fig 4B). Poly I:C activation of the intracellular receptor TLR3 induced strong phosphorylation of signaling kinases, TBK1, ERK1/2 as well as activation of Akt and p65 which were also disrupted in EIPA treated cells (Fig 4C). Interestingly, PAM3CSK4 activation of TLR1/2 generated signaling at the cell surface which was not disrupted by EIPA inhibition of macropinocytosis (Fig 4D). These findings are consistent with the well-entrenched reputation of TLR9 as an endosomal receptor, however, the dependence on EIPA suggests that macropinosomes are an essential site and/or portal for TLR9 activation and downstream signaling.

### Acidification of macropinosomes as a TLR9 signaling environment

Other studies have concluded that TLR9 signals from lysosomes, based on the requirements for an acidic and degradative environment for TLR9 ecto-domain proteolytic cleavage, a necessity for signaling through the MyD88 adaptor (Ewald et al., 2008, Park et al., 2008). It is relevant that macropinosomes are also known to be acidified (Canton, 2018, Swanson, 2008) and known to contain cathepsins which are needed, notably, for viral entry (Izumida, Hayashi et al., 2020, Krzyzaniak, Zumstein et al., 2013). To assess these characteristics, BV-2 cells were incubated with the fluorescent tracer DQ-BSA, which fluoresces upon acid-dependent proteolysis (Fig 5A, Appendix Fig S5A). By bathing cells in TMR-dextran (red) and DQ-BSA, we recorded newly formed macropinosomes that contained both dextran and fluorescing (green) DQ-BSA. The live imaging and quantification indicate that DQ-BSA fluorescence increases steadily within the first 6 mins, at a time when the marker should be exclusively in macropinosomes. DQ-BSA also fluoresced in other compartments in line with the acidic, degradative nature of later endolysosomal environments (Appendix Fig S5B).

**Figure 5.**
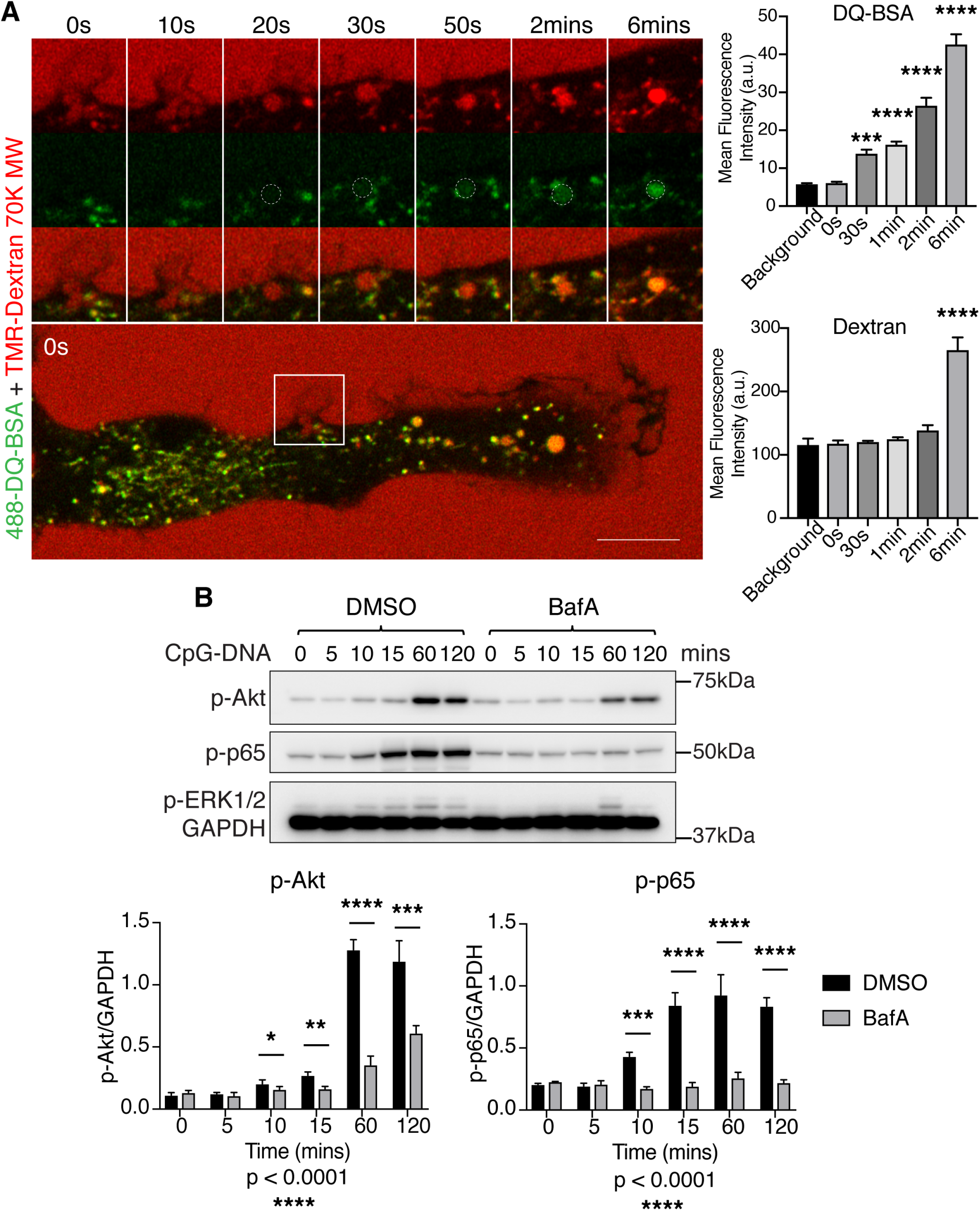
Rapid macropinosome acidification supports proteolytic cleavage and TLR9 signaling. **A**. Live cell confocal imaging of BV-2 microglia incubated with 488-DQ-BSA (green) and TMR-Dextran (red). Movie sequence follows an individual macropinosome from formation to maturation. Mean DQ-BSA and dextran fluorescence intensity within macropinosomes are measured over time (ordinary one-way ANOVA with Dunnett’s multiple comparisons test, significance between control and fluorescence intensity across each time point displayed on graph). **B**. Immunoblot and quantification of BV-2 microglia untreated and treated with Bafilomycin A1 (BafA) and activated with CpG-DNA over a 2 hour time course (two-way ANOVA with Sidak’s multiple comparisons test, significance for interaction marked underneath graph, significance between DMSO and BafA treated samples at specific time points are marked on graph). Data information: Graphs are displayed as mean ± SD. n=3 independent experiments for all panels.

The need for an acidic environment for TLR9-induced MyD88-dependent signaling was further tested in cells treated with an inhibitor of the vacuolar-type proton-ATPase, Bafilomycin A1 (BafA), which dissipates acidification throughout endocytic/macropinocytic pathways (Yoshimori, Yamamoto et al., 1991). The activation of p65 was assessed as an example of MyD88 dependent signaling and phospho-p65 induced by CpG-DNA from 10 min and throughout a 2 hour period was negated by BafA (Fig 5B). Akt phosphorylation which is initiated in macropinosomes was also disrupted but there was some recovery at later time points. BafA ablated both early and late activation of p65 indicating that acidification does contribute to TLR9/MyD88 signaling (Fig 5B). Although not quantified, phospho-ERK1/2 was also seen to be ablated, particularly at early times by BafA.

Taken together the DQ-BSA staining and the ablation of signaling by BafA indicate that macropinosomes can provide an acidified and proteolytic environment for TLR9 signaling, including through MyD88.

### In microglia, macropinosomes host TLR-activated LRP1 and Rab8a

Cell turnover, injury and death contribute proteinaceous debris and nucleic acids to the brain environment during growth and ageing (Mandavilli, Santos et al., 2002, Von Bernhardi, Eugenín-von Bernhardi et al., 2015, Yankner, Lu et al., 2008). Cell debris and nucleic acids can be internalized by macropinocytosis, as demonstrated by DNA staining in macropinosomes of primary microglia (Appendix Fig S6). With TLR9 and CpG also present in these compartments, it is essential to have mechanisms in place to constrain pro-inflammatory responses which might otherwise be chronically activated in these scavenging cells. In macrophages, macropinosomes host a regulatory complex composed of LRP1/Rab8a and PI3Kγ that biases TLR-induced signaling and inflammatory cytokines towards an anti-inflammatory axis (Luo et al., 2018, Luo et al., 2014). We thus investigated whether the LRP1/Rab8a/PI3Kγ complex is present in microglia and whether it influences TLR9 signaling and outputs.

tdTomato-Rab8a decorates 488-dextran filled macropinosomes at the periphery of BV-2 cells, and imaging reveals that the GTPase is recruited to the membranes just at the point of macropinosome formation on ruffles (Fig 6A). Co-expression of GFP-Rab8a with LifeAct-Scarlett as an F-actin probe, confirm that Rab8a is on macropinosomes preceding their closure and it is retained for a short while after depolymerization of the actin (Fig 6B). Thus, Rab8a is recruited spatially and temporally in similar fashion in macrophages (Wall et al., 2019, Wall et al., 2017) and microglia as a transient macropinosome resident.

**Figure 6.**
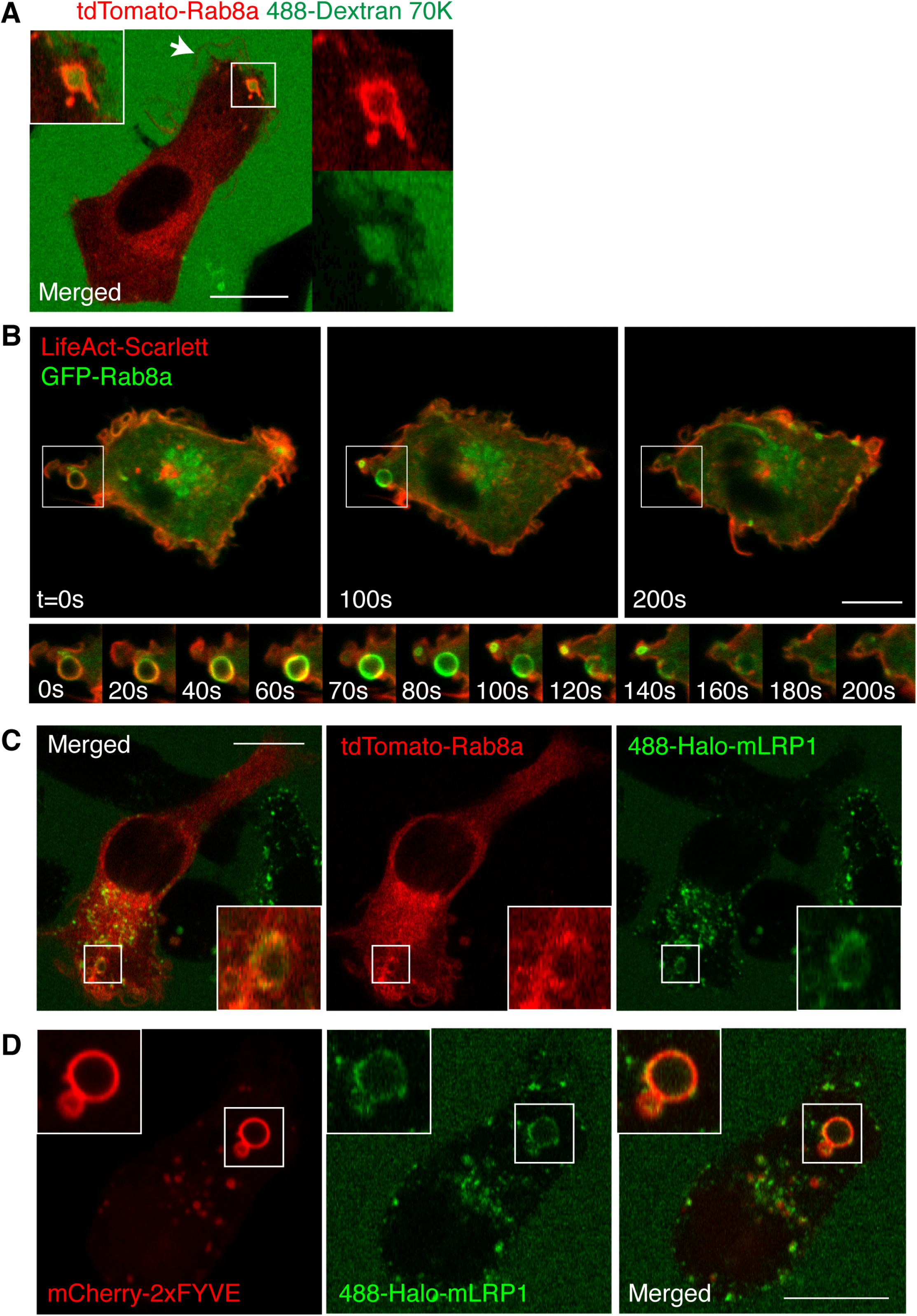
LRP1 and Rab8a are localized to the macropinosome. **A**. Single confocal slice in time of live BV-2 microglia transiently transfected with tdTomato-Rab8a incubated with 488-dextran. **B**. Movie sequence following BV-2 cell transiently transfected with GFP-Rab8a and LifeAct-Scarlett. Inset focuses on formation and maturation of a single macropinosome. **C**. Single slice in time from live cell imaging of tdTomato-Rab8a transiently transfected in Halo-mLRP1 stably expressing BV-2 microglia. **D**. Single time slice from live cell imaging of 2XFYVE transiently transfected in Halo-mLRP1 stably expressing BV-2 microglia. Data information: All imaging representative of population. All scale bars represent 10μm.

LRP1 is an abundant endocytic receptor that recycles constantly and is present at the cell surface, in recycling endosomes and in multiple endocytic pathways as reflected by immunostaining in microglial cells (Appendix Fig S7). LRP1 transits briefly through macropinosomes before it is sorted for recycling or degradation. Establishing the joint presence of LRP1and Rab8a in microglial macropinosomes involved the expression of an LRP1 mini-receptor construct (mLRP1) used previously for trafficking studies (Li, Marzolo et al., 2000), to which we attached an exofacial HaloTag, termed Halo-mLRP1 (labelled with 488-Halo ligand) (Appendix Fig S8). 488-Halo-mLRP1 co-expressed in BV-2 cells produced punctate labeling that included puncta around tdTomato-Rab8a-labeled macropinosomes (Fig 6C). Some 488-Halo-mLRP1 was also found on macropinosomes, after loss of Rab8a and maturation of the compartments, now demarked with the PI3P marker, mCherry-2xFYVE (Fig 6D). Thus, key elements of the anti-inflammatory complex, LRP1 and Rab8a, define early stages of macropinosomes in BV-2.

Showing that LRP1 and Rab8a form a complex in microglia, entailed first, immunoprecipitation of LRP1 from BV-2 cells extracted after incubation with or without CpG-DNA. CpG-DNA induced phosphorylation of LRP1 indicated by phospho-tyrosine (p-Tyr) detection of immunoprecipitated LRP1 and Rab8a was found to co-precipitate with LRP1, primarily after activation (Fig 7A). To establish that TLR-induced activation of LRP1 and its interaction with Rab8a occur during macropinocytosis, cells were exposed to CpG-DNA and other TLR ligands and treated with the inhibitor EIPA prior to LRP1 immunoprecipitation (Fig 7B). The phosphorylation of LRP1 induced by LPS, Poly I:C or CpG-DNA was decreased in EIPA treated cells. Similarly, the co-immunoprecipitation of Rab8a was also decreased after inhibition of macropinocytosis, markedly in the case of CpG-DNA and Poly I:C activated cells. Thus, LRP1 and Rab8a form a complex and both are activated downstream of TLR9 in the context of early macropinosomes in microglia. Next, the activation of Rab8a by CpG-DNA was demonstrated using capture of GTP-Rab8a on a Rab binding domain probe from an effector, the inositol polyphosphate 5-phosphatase, OCRL (OCRL-RBD). Pull down of active GTP-bound Rab8a using GST-OCRL beads demonstrated that Rab8a activation is significantly induced by CpG-DNA with a peak observed at 15-30 mins (Fig 7C). Finally, the capture of the effector PI3Kγ by active Rab8a can be demonstrated by loading Rab8a in BV-2 lysates with a non-hydrolysable GTP (Gpp(NH)p). GTP-Rab8a preferentially pulled down the p110 subunit of PI3Kγ (Fig 4D). Taken together, these data reveal in microglia that, downstream of TLR9 and in a macropinocytosis-dependent manner, LRP1 and Rab8a are activated and co-recruited, serving to further recruit the effector PI3Kγ.

**Figure 7.**
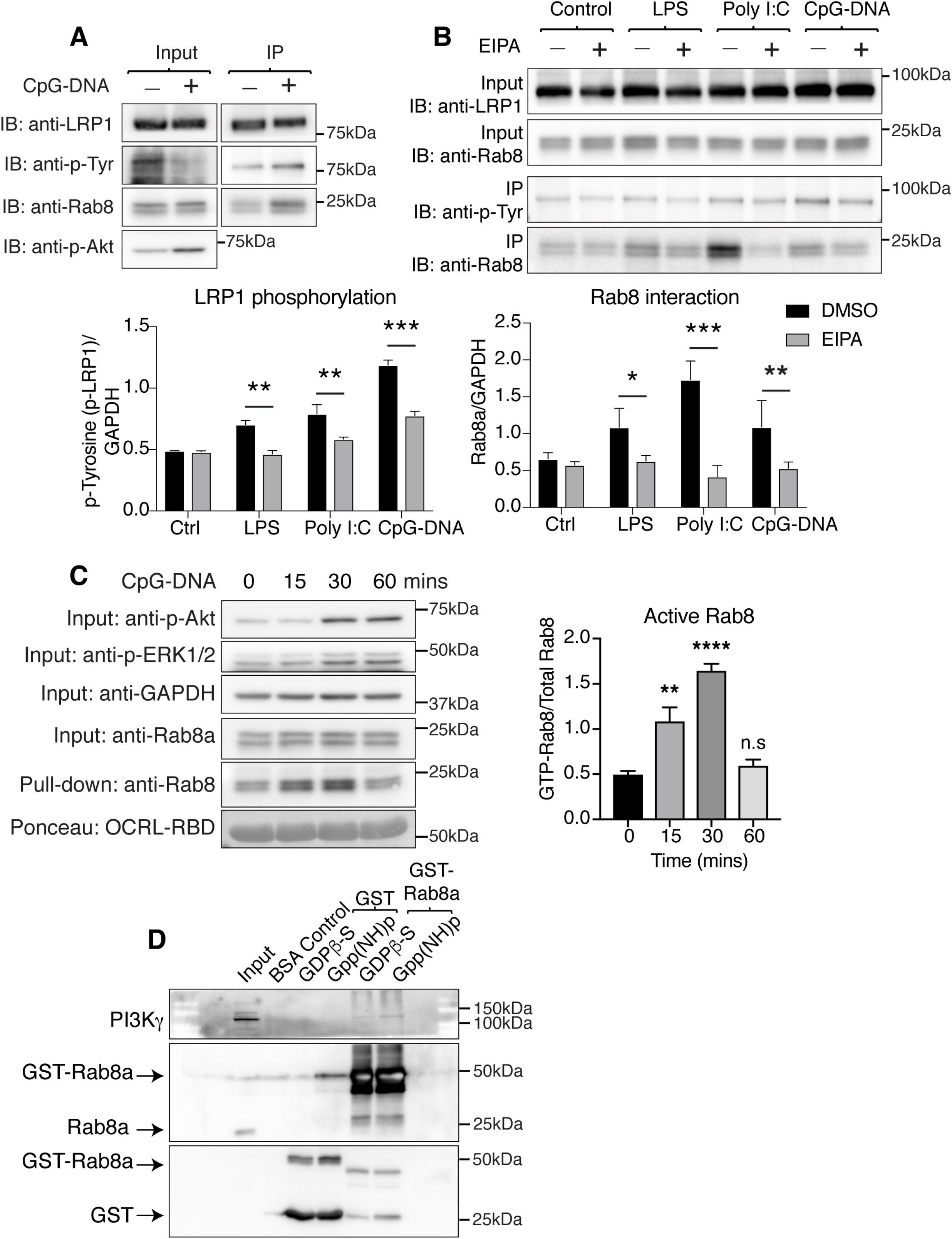
CpG-DNA activation, interaction of LRP1/Rab8a during macropinocytosis and GTP-Rab8a capture of PI3Kγ. **A**. Immunoprecipitation of LRP1 using an anti-LRP1 antibody. BV-2 cells were incubated with or without CpG-DNA (60 mins) prior to extraction. Immunoblotting of lysates p-Akt confirms activation of the cells and LRP1 phosphorylation in the IP fraction is detected by a phospho-tyrosine (p-Tyr) antibody. Rab8a (upper band of Rab8 doublet) is co-immunoprecipitated with LRP1 after CpG-DNA activation. **B**. TLR-induced LRP1 activation and Rab8a binding. Lysates were prepared from BV-2 microglia pre-treated or not with EIPA and activated with different TLR ligands (LPS, Poly I:C, CpG-DNA). Anti-phospho-Tyrosine (p-Tyr) and anti-Rab8a blotting were used to detect LRP1 phosphorylation and Rab8a coprecipitation. Representative gel lanes shown and data from 3 independent experiments was used to quantify ligand induction and EIPA attenuation of LRP1 phosphorylation and Rab8a binding, respectively. \*\*\*\**p<0*.*0001* for both LRP1 phosphorylation and Rab8a interaction (two-way ANOVA with Sidak’s multiple comparisons test, significance marked on graph for comparison across drug treatment). **C**. CpG-DNA activation of (GTP-bound) Rab8. Phospho-Akt and phospho-ERK1/2 blotting in cell lysates demonstrates activation of BV-2 cells by CpG-DNA over time. GAPDH and Rab8a blotting in the input samples are loading controls. A probe based on the Rab binding domain of the effector OCRL (OCRL-RBD) was used to pull down GTP-Rab8 from lysates. Ponceau stain demonstrates even loading of the probe. CpG-DNA induced GTP-Rab8 is measured by densitometry and displayed in graph (ordinary one-way ANOVA with Dunnett’s multiple comparison’s test, significance between control and time course displayed on graph). **D**. Pull down of PI3Kγ by Rab8. BV-2 cell lysates were incubated with GST-Rab8a beads in medium containing non-hydrolysable GDP (GDPβS) or non-hydrolysable GTP (Gpp(NH)p). The effector PI3Kγ (p110 catalytic subunit) be detectably captured on GST-Rab8a beads after GTP-loading and less so with GDP-loaded beads. BSA and GST controls demonstrate the absence of non-specific binding. Rab8a blotting captures Rab8a in the input and on GST-Rab8a beads. GST blotting detects GST-only and GST-Rab8a beads. Data information: All blots are representative. Graphs are displayed as mean ± SD. For statistical analysis, n=3 independent experiments.

### Constraining TLR9-induced pro-inflammatory responses

Functional evidence that LRP1/Rab8a, with PI3Kγ as an effector, modulate TLR9-induced inflammatory responses, was next addressed by examining signaling and cytokine production. BV-2 cells pre-treated with the PI3Kγ isoform-selective inhibitor IPI-549 prior to activation with CpG-DNA were assessed in replicate experiments for the activation of PI3K/Akt signaling. Phospho-Akt levels as well as other PI3K/Akt/mTOR associated signaling proteins and Akt substrates (phospho-PRAS40, phospho-mTOR, phospho-p70S6K) were all activated by CpG-DNA and found to be markedly depressed by IPI-549 inhibition (Fig 8A). Phospho-p65, as part of the NF-kB signaling pathway, and as a direct outcome of MyD88 recruitment, saw no change downstream of TLR9 in cells treated with IPI-549 and there was only a modest effect on phospho-ERK1/2 levels in these cells. Despite being dependent on macropinocytosis (see Fig 4) the activation of p65 and ERK1/2 are not influenced by PI3Kγ. Phospho-Akt expression was measured and found to be significantly decreased upon IPI-549 treatment, consistent with a dependence on PI3Kγ.

**Figure 8.**
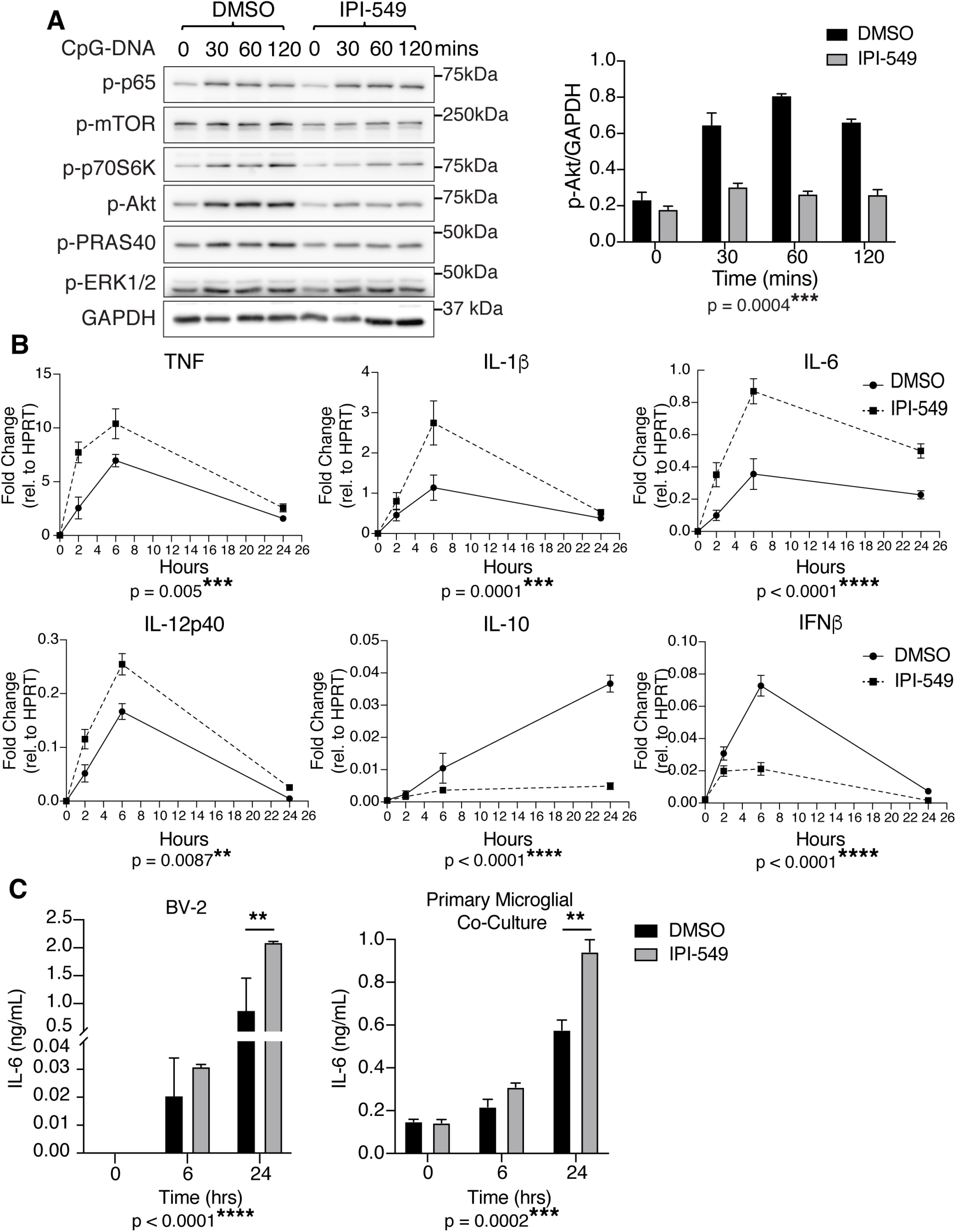
PI3Kγ is required for TLR9-driven Akt/mTOR signaling and delimits the production of proinflammatory cytokines. **A**. Immunoblotting of BV-2 cells untreated and treated with selective PI3Kγ inhibitor, IPI-549 and activated over a CpG-DNA time course (0-120 mins). Expression levels of phospho-Akt (p-Akt) was quantified relative to control, GAPDH by densitometry. **B**. Expression of proinflammatory and anti-inflammatory cytokine genes from BV-2 microglia untreated and treated with IPI-549 and activated with CpG-DNA across a time course of 24 hours. n=3 independent experiments (two-way ANOVA with Sidak’s test for multiple comparisons, significance for interaction displayed underneath graph). **C**. IL-6 secretion from microglia cell line BV-2 (left) and primary microglial co-cultures (right) treated with DMSO control versus IPI-549 and activated with CpG-DNA over 24 hours. Interaction was significant for BV-2 at \*\*\*\**p<0*.*0001 (t=24, p=0*.*0013)*. Interaction was also significant for primary microglial co-culture with \*\*\**p = 0*.*0002 (t=24, p=0*.*0017)*, (two-way ANOVA with Sidak’s test for multiple comparisons). BV-2: n=3 independent experiments, primary co-culture: n=5 animals. Data information: Graphs are displayed as mean ± SD.

Inhibition of PI3Kγ by IPI-549 also resulted in a pronounced pro-inflammatory status, shown through qPCR results that demonstrate increased expression of pro-inflammatory cytokines TNF, IL-1β, IL-6 and IL-12p40 across a 24 hour CpG-DNA activation time course (Fig 8B). Correspondingly, IPI-549 treatment resulted in a decrease in the production of anti-inflammatory mediators, IL-10 and IFNβ expression (Fig 8B). The secretion of IL-6 was assessed as a prominent pro-inflammatory cytokine downstream of TLR9 in BV-2 cells and in primary microglial co-cultures. IPI-549 treatment significantly increased IL-6 secretion compared to DMSO controls in both BV-2 and primary co-culture (Fig 8C). We can conclude based on these results that, in microglia, PI3Kγ normally acts to constrain inflammation through Akt signaling downstream of TLR9 and through the biasing of cytokine programs towards anti-inflammatory mediators.

### PI3Kγ recruits Akt for TLR9 signaling in macropinosomes

Early macropinosomes are equipped as signaling hubs through the PI3K conversion of PI(4,5)P_2_ to PI(3,4,5)P_3_ which is transiently enriched in newly-closed macropinosomes and acts as a platform for recruiting signaling kinases including Akt (Swanson & Yoshida, 2019, Yoshida et al., 2009). Live cell imaging of a fluorescently tagged Akt1 reporter (TagRFP-Akt1) (Norris, Yang et al., 2017) was used to detect Akt recruitment to signaling membranes in CpG-DNA treated BV-2 cells (Fig 9A). In control cells, TagRFP-Akt1 was largely cytosolic and enriched at cell surface ruffles, presumably at the behest of ongoing, serum-induced receptor activation. Strikingly, after addition of CpG-DNA, TagRFP-Akt1 was now found concentrated on macropinosomes (Fig 9A). In activated BV-2 cells, TagRFP-Akt1 and GFP-Rab8a were both recruited to ruffles and newly forming macropinosomes. Notably, the Akt reporter was not present on Rab8a sorting tubules emerging from the macropinosomes (Fig 9B). This is direct evidence for early macropinosomes as a signaling site for TLR9-induced Akt recruitment.

**Figure 9.**
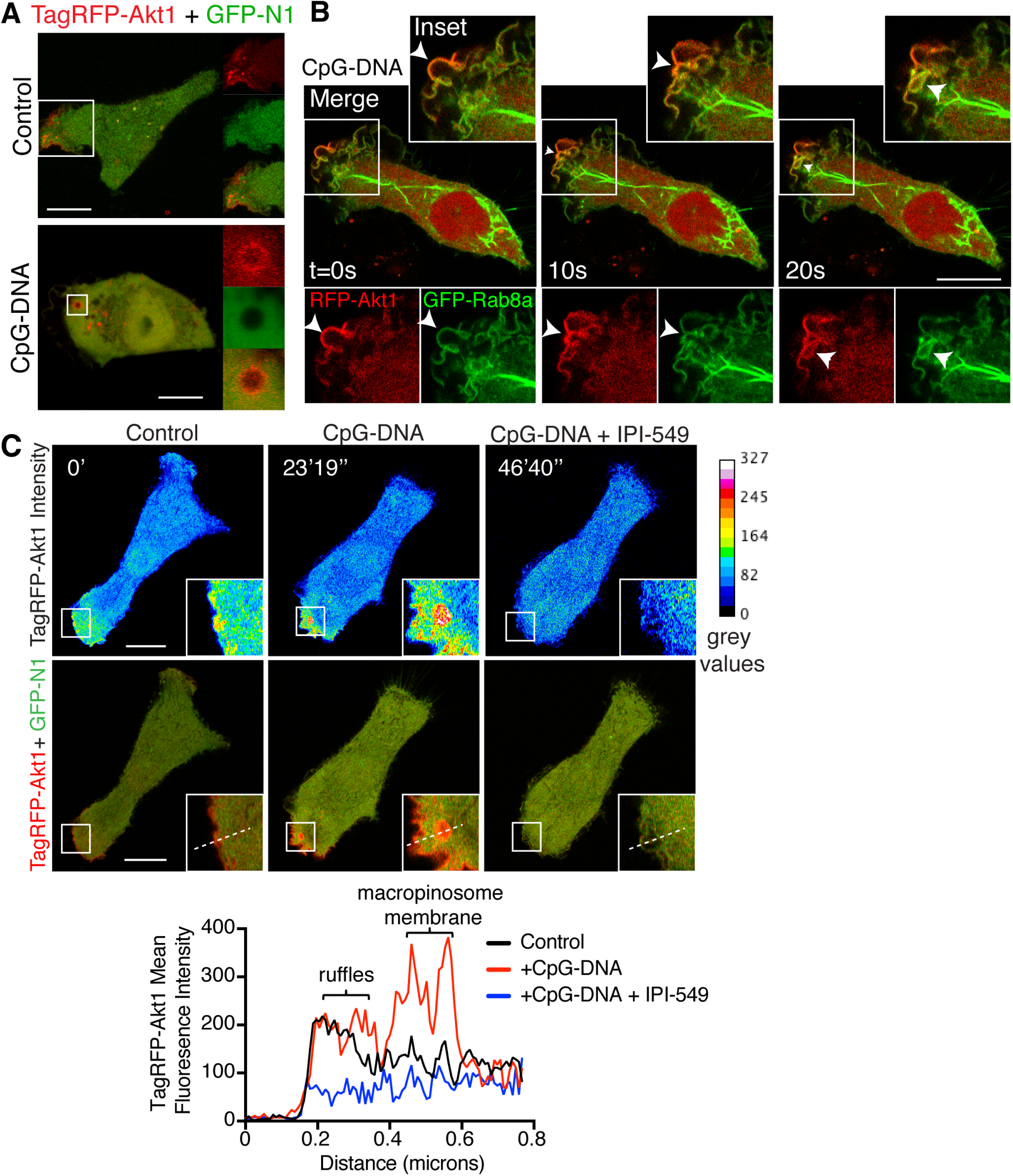
Akt is recruited to TLR9-signaling macropinosomes in a PI3Kγ-dependent manner. **A**. Single confocal image in time from live cell imaging of the Akt reporter, TagRFP-Akt1 (red) and ratiometric control, GFP-N1 (green) transiently co-transfected in unstimulated and CpG-DNA activated BV-2 microglia. **B**. Live confocal imaging of GFP-Rab8a and TagRFP-Akt1 transiently expressed in BV-2 microglia demonstrate co-localisation on ruffling membranes and forming macropinosomes (white arrowhead). **C**. Live cell imaging of a single BV-2 cell transiently transfected with TagRFP-Akt1 (red) and a cytosolic GFP-N1 (green) as a ratiometric control. TagRFP-Akt1 intensity is also displayed as a heatmap to highlight changes in intensity. The imaging captures TagRFP-Akt1 at rest (unstimulated, left panel) and recruitment and enrichment on macropinosomes after activation with CpG-DNA (CpG-DNA, center panel). This fluorescence is then ablated upon addition of the PI3Kγ-selective inhibitor, IPI-549 (CpG-DNA+IPI-54). Fluorescence of TagRFP-Akt is measured as line scans across the same region encompassing the cell edge and ruffling/macropinocytic domains at different times. All events are captured in the one cell. Data information: Graphs are displayed as mean ± SD. All scale bars represent 10μm.

Finally, we implicate PI3Kγ in Akt recruitment using sequential imaging of BV-2 cells to compare CpG-DNA-induced recruitment of TagRFP-Akt1 before and after addition of CpG-DNA and the PI3Kγ-specific inhibitor IPI-549. TagRFP-Akt1 is observed in a representative live cell on the plasma membrane and ruffling edge, as depicted by line scan measurements of mean TagRFP-Akt1 fluorescence intensity (Fig 9C). Upon addition of CpG-DNA to the imaging medium, stronger enrichment of TagRFP-Akt1 is observed in new clusters on surface ruffles and it appears on newly formed macropinosomes which line scans show is the strongest site of recruitment (Fig 9C). Upon addition of the IPI-549 inhibitor, the TagRFP-Akt1 fluorescence disappears from the ruffles and from any newly forming macropinosomes (one of which is shown in the inset) (Fig 9C). The CpG-induced recruitment of the Akt probe to forming macropinosomes is PI3Kγ dependent. The broader loss of Akt fluorescence at the ruffle in response to IPI-549 may indicate that PI3Kγ is the predominant PI3K acting in CpG-DNA activated cells. These results overall depict Akt recruitment to ruffles and new macropinosomes as a result of TLR9 activation and show a role for PI3Kγ in TLR9-induced signaling.

## Discussion

With the ability to launch inflammatory and immune responses, TLRs are important for the customized protective and regenerative roles of microglia (Wolf et al., 2017). In this study we characterized inflammatory responses downstream of the intracellular receptor TLR9, which is poised to detect microbial and host nucleic acids earmarked by unmethylated CpG motifs, both of which are highly relevant to the scavenging role of microglia in CNS environments. Macropinocytosis is a prominent pathway for ingestion of fluid, cell debris and pathogens (Canton, 2018, Mercer & Helenius, 2009) and we show in primary microglia that this pathway is significantly upregulated by TLR9 activation. The prominence of macropinosomes as a capacious compartment for fluid-phase and membrane internalization and for directing this cargo through endolysosomal pathways to degradative compartments, cannot be ignored in the context of TLR activation and signaling. Indeed, our live cell imaging reveals TLR9 and exogenous CpG-DNA are located in macropinosomes, in addition to other endolysosomes. These macropinosomes are acidified, proteolytic sites that do support and are required for CpG-DNA activated TLR9 signaling. The physiological importance of macropinosomes in this capacity is highlighted by access to macropinosomal signaling regulators, including LRP1 and PI3Kγ that constrain inflammation triggered by TLR9, with the potential to mitigate inflammatory damage in the CNS.

In microglia, recombinant TLR9 localized in macropinosomes, lysosomes and other endocytic compartments, with little of the receptor found on the cell surface or in the ER. In dendritic cells, TLR9 is localized mainly in the endoplasmic reticulum (ER) (Leifer et al., 2004), while this is not necessarily the case in other cell types, such as B cells (Chaturvedi et al., 2008, Chaturvedi & Pierce, 2009). The trafficking of TLR9 was initially shown to be direct from the ER to lysosomes (Kim, Brinkmann et al., 2008, Latz et al., 2004, Park et al., 2008) and central to this transport for endosomal TLRs is the transmembrane chaperone UNC93B1 (Kim et al., 2008). TLR9 transport from the ER to lysosomes via the Golgi has also been reported (Chockalingam, Brooks et al., 2009). UNC93B1 was subsequently shown with additional roles in endosomal TLR trafficking, including packaging TLR9 into COPII vesicle for transport from the ER to the Golgi, and thence for further post-Golgi trafficking (Lee et al., 2013). UNC93B1 recruits the trafficking adaptor AP-2 to shepherd TLR9 internalization through clathrin-mediated pathways, as an alternative and selective entry point for the TLR9, but not for TLR7, into the endolysosomal pathway (Lee et al., 2013). While this finding established surface delivery and internalization of TLR9, others have speculated and our data now confirm, that macropinocytosis is a more likely and abundant pathway for TLR9 internalization from the surface and for its ultimate delivery to lysosomes through macropinosomal maturation.

Macropinocytosis is also a major pathway for the entry or scavenging of viral and bacterial pathogens and therefore a likely route for TLR9 engagement (Mercer & Helenius, 2009). The identification of macropinosomes as the TLR9 uptake vacuoles is based on defining characteristics presented here, such as recording their formation from ruffles in live cell imaging, dextran uptake, Rab8a labeling (Wall et al., 2019) and their inhibition by EIPA. Interestingly imaging of 647-CpG-DNA uptake showed that in addition to fluid-phase entry, membrane bound CpG-DNA buds off early macropinosomes in what could reflect secondary clathrin-mediated uptake from macropinosomes, neatly consolidating these pathways. Notably, the prevalent recycling receptor and clathrin-coated vesicle cargo, LRP1, was tracked into macropinosomes where it appears as very similar puncta or vesicles around the macropinosomes in microglia (Fig 5C, D).

A major revelation from this study is that ligand contact and signaling for TLR9 occur in macropinosomes, much earlier in endocytic pathways than previously reported. The requirements for acidification for TLR9 ectodomain cleavage, along with a degradative environment to expose nucleic acids, have previously focused attention on lysosomes as the major signaling site for this receptor (Ahmad-Nejad et al., 2002, Häcker et al., 1998, Latz et al., 2004, Park et al., 2008). However, macropinosomes also ostensibly fulfill the requirements for TLR9 modification since they too are acidified (Canton, 2018) and support proteolytic cleavage of some ligands and receptors as prerequisites for their sorting or signaling in this compartment (Krzyzaniak et al., 2013). Such features are directly evidenced here by the coincidence of acid and cleavage-dependent DQ-BSA fluorescence along with dextran uptake in early macropinosomes and by bafilomycin inhibition of TLR9 and MyD88-dependent p65 phosphorylation. Preventing acidification of the macropinosomes had a significant impact on TLR9 signaling in our hands and similar experiments have also been done for late stage TLR9 signaling (1-2 hours post CpG-DNA treatment) within lysosomes (Combes et al., 2017, Häcker et al., 1998, Latz et al., 2004).

CpG-DNA uptake was halved but nearly all signaling blocked when macropinocytosis was inhibited by EIPA, implying first, that CpG-DNA entering cells through macropinocytosis is responsible for triggering TLR9, and second, that macropinosomes themselves or downstream compartments are TLR9 signaling sites. Our results do not exclude TLR9 signaling from also occurring in endosomes or lysosomes downstream of macropinosomes and TLR9-induced signaling continued for some time beyond 5-15 mins, peaking at 60 mins post activation, which is in line with previous reports (Ahmad-Nejad et al., 2002, Häcker et al., 1998, Latz et al., 2004). Phosphorylation of Akt and downstream p65 are established here as being EIPA sensitive and thus macropinocytosis dependent, moreover this signaling profile resulted in both pro-inflammatory and as well as IFNβ cytokine production, all of appear to be driven from the same compartment. Early macropinosomes are equipped for signaling through the transient enrichment of PI3K-generated PI(3,4,5)P_3_ (Swanson & Yoshida, 2019, Wall et al., 2017) as a platform for recruiting Akt and other signaling kinases, demonstrated here directly by imaging the recruitment of an Akt reporter. TLR9 recruits MyD88 as its only traditional adaptor and our results are consistent with this signaling being generated from macropinosomes. In other cell types, there is evidence for TLR9 signaling from different endolysosomal compartments, in response to different types of CpG-DNA, which also differentiates the production of pro-inflammatory cytokines and type I interferon responses (Lee & Barton, 2014). In macrophages, a variety of stress inducers can enhance macropinosome fusion with lysosomes, phagosomes or autophagosomes (Racoosin & Swanson, 1993, Yoshida, Pacitto et al., 2015) and this could alter the signaling environment for TLR9. In the microglia used here, there was no evidence for stress-induced fusion of compartments since Lamp1 staining was not seen in early macropinosomes, nor was there swelling of endolysosomal compartments.

CpG-DNA capture by ruffles and non-selective entry into macropinosomes highlights the potential for unfettered exposure of TLR9 in these compartments to any microbial or host nucleic acids in the cell or tissue milieu. The sequestration of TLR9 in lysosomes for signaling has been proposed as a key mechanism for avoiding activation of TLR9 by host DNA, which can otherwise lead to excessive inflammation and autoimmunity (Barton et al., 2006). Engineering the cell surface placement of signaling-competent TLR9 in cells or transgenic mice starkly highlights the potential for environmental, host nucleic acid to trigger hyper- and chronic-inflammation (Barton et al., 2006, Mouchess, Arpaia et al., 2011) and even neonatal fatality (Stanbery, Newman et al., 2020). While, the presence of TLR9 and its signaling capacity in macropinosomes might elevate surveillance and defense capabilities for microglia, this may happen at the expense of promiscuous and even chronic TLR9 activation. The transient nature of the signaling domains in this compartment and the rapid sorting of macropinosome contents might help to limit inflammatory responses. The presence of mitigating signaling regulators in macropinosomes, described below, is also of prime importance. However, the ability of TLR9 to be activated and signal in macropinosomes now prompts further deliberation about how aberrant or self-DNA triggered TLR9 activation is avoided, since macropinosomes do not offer the same level of sequestration as lysosomes.

LRP1 is a multi-functional endocytic receptor and one of its roles is as an anti-inflammatory regulator in TLR pathways and in other settings including atherosclerosis (Xian, Ding et al., 2017) and neurodegenerative disease (Brifault, Gilder et al., 2017, Yang, Liu et al., 2016). LRP1 is activated as a TLR4 co-receptor for recruitment of Rab8a in macrophages (Luo et al., 2018), and the results here show that these regulators are similarly activated by TLR9 and recruited in microglia. The LRP1/Rab8a complex can then recruit PI3Kγ to augment Akt and mTOR signaling downstream of activated TLR9. The spatiotemporal recruitment of PI3Kγ to early macropinosomes and the effect of its kinase activity are now graphically demonstrated by live imaging of CpG-DNA-induced Akt recruitment to these membrane domains in live cells. Through this Akt signaling, PI3Kγ biases the cytokine profile, downregulating pro-inflammatory cytokines and upregulating anti-inflammatory mediators in microglia, consistent with similar effectors in mouse and human macrophages demonstrated with genetic ablation or pharmacologic inhibition of PI3Kγ (Kaneda et al., 2016, Luo et al., 2014, Wall et al., 2017).

In microglia, PI3Kγ is highly expressed and upregulated by LPS where it can function through kinase dependent and independent roles (Frister, 2014, Lajqi, Lang et al., 2019). In other microglial pathologies, PI3Kγ can be neuroprotective by restraining microglial toxicity, which is in line with our findings, after focal brain ischemia (Schmidt, Frahm et al., 2016), but it can also drive neuroinflammation after surgical brain injury in rats (Huang, Sherchan et al., 2015). In other cell and *in vivo* scenarios, PI3Kγ is known to be active in TLR9 pathways including in CpG-induced pleurisy (Lima, Marques et al., 2019) and downstream of mtDNA activated TLR9 in models of cardiotoxicity (Li, Sala et al., 2018). The specific role of PI3Kγ downstream of TLR9 in microglia is addressed here for the first time and now requires further elucidation given the translational value of its pharmacological inhibitors for inflammation and disease.

In summary, we present the macropinosome as an early signaling compartment for TLR9 which facilitates the uptake of CpG-DNA and supports MyD88-dependent signaling. This places TLR9 at the frontline of encountering host and microbial nucleic acids for activating innate immune and inflammatory responses in microglia. In this environment, inflammation triggered by TLR9 can be modulated by regulators such as LRP1 and PI3Kγ to potentially mitigate hyperinflammation.

## Materials and Methods

### Antibodies and reagents

All antibodies and reagents used for experiments are listed in the Reagents and Tools table.

#### Reagents and Tools Table

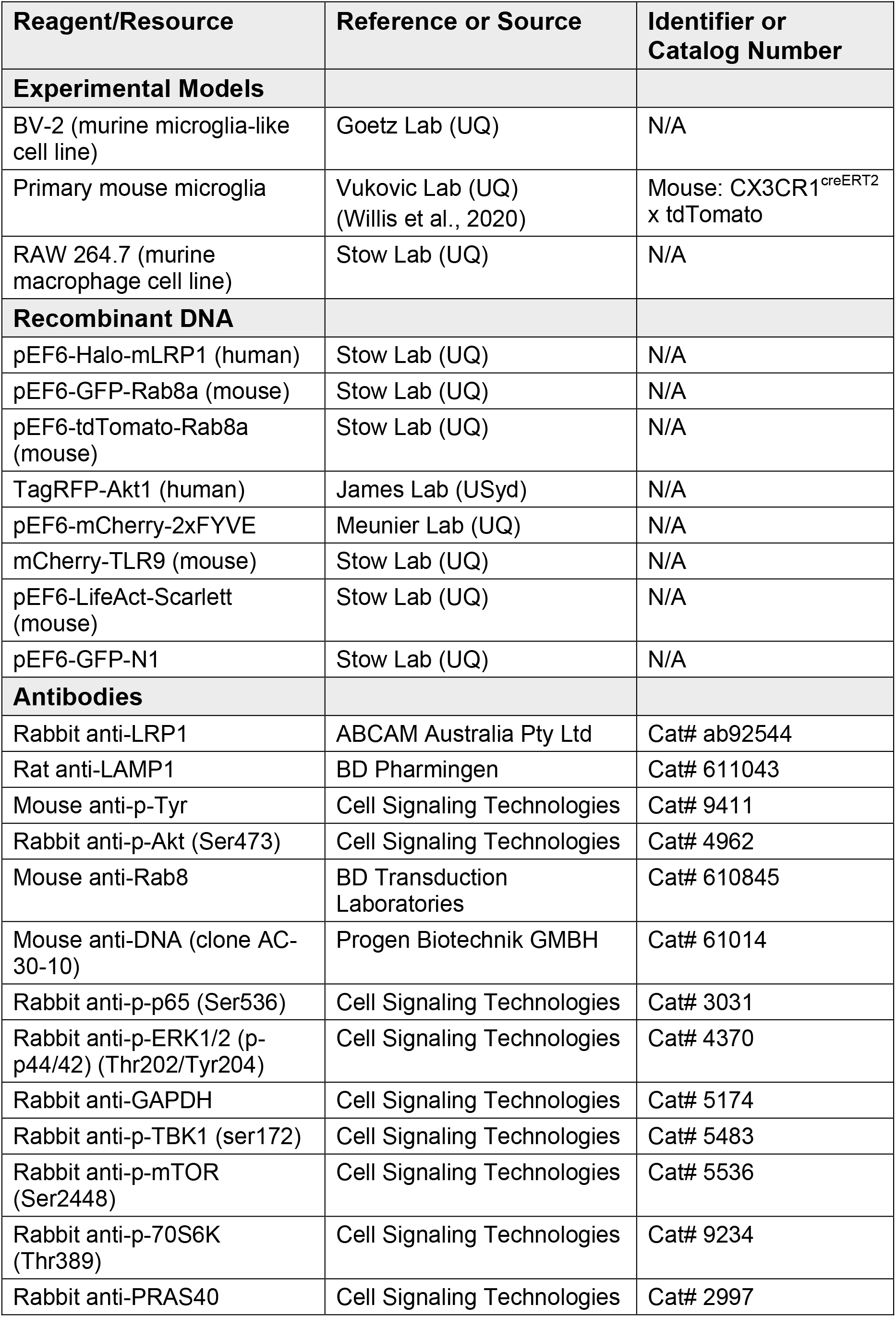
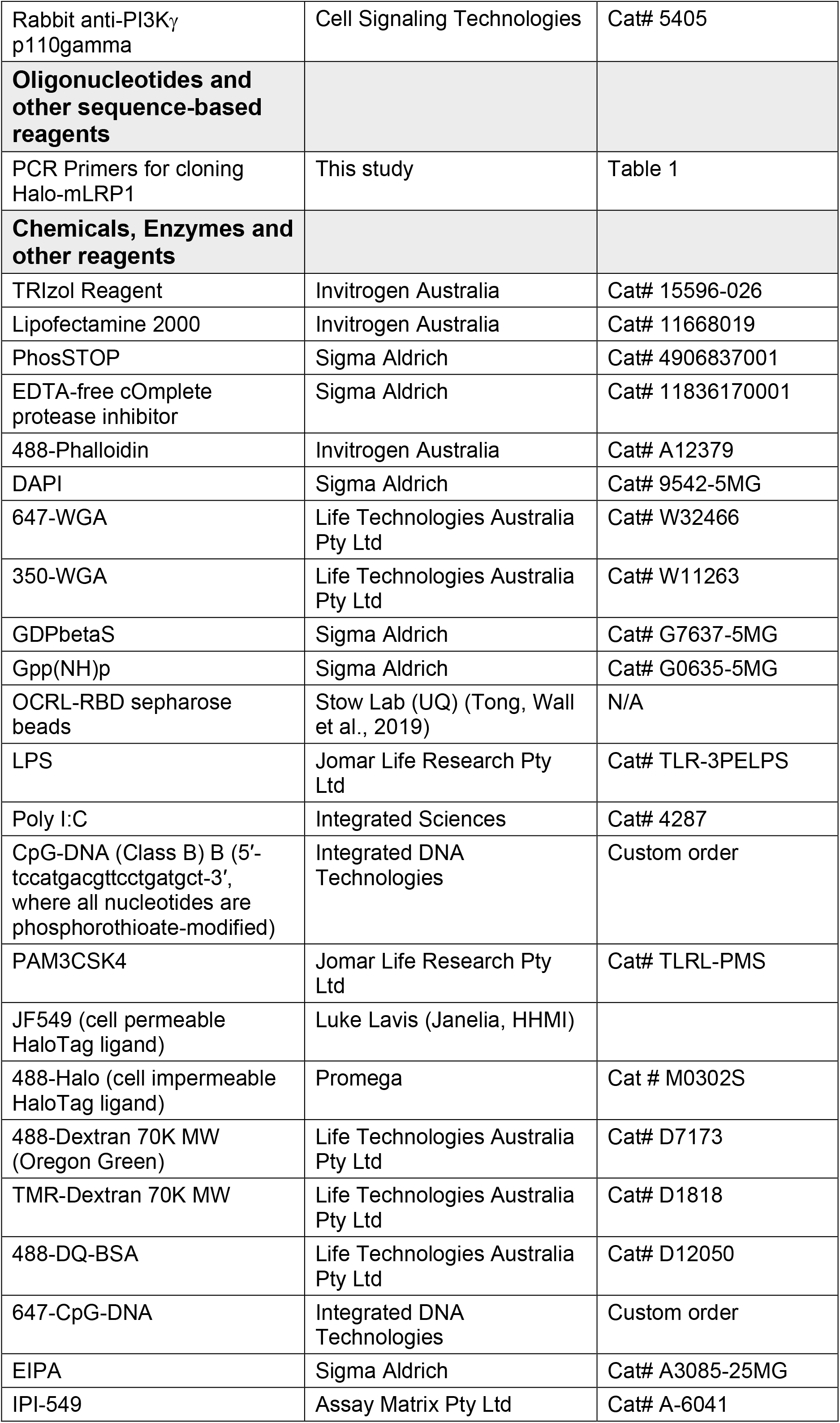
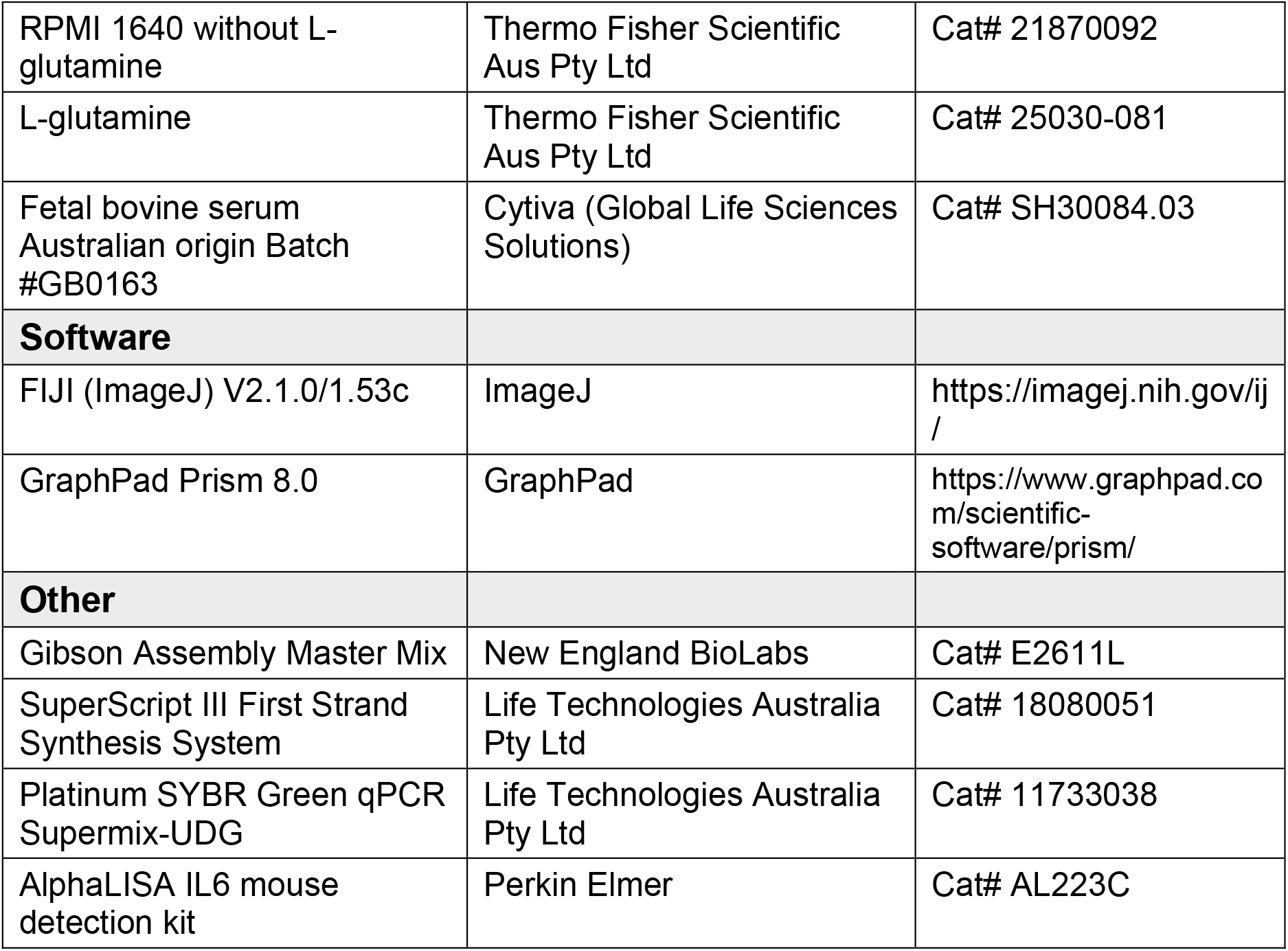

### Cloning and constructs

All plasmids and constructs used for experiments are listed in the Reagents and Tools table. The Halo-tagged LRP1 minireceptor (Halo-mLRP1) was developed through restriction digest (NheI and AsiSI) and ligation of HaloTag from Halo-Rab13 (Condon et al., 2018) and Gibson Assembly as per manufacturer’s instruction (NEBiolabs) of a gblock (IDT Technologies) containing 2XMyc and a LRP1 signal peptide (Table 1) into an empty pEF6-V5-HIS-TOPO backbone. The minireceptor LRP1 sequence was amplified by PCR using the mLRP4T100 minireceptor construct as a template (primers described in Table 1). The mLRP1 fragment was then purified and inserted into the pEF6 backbone. Aside from transient expression by transfection, Halo-mLRP1 was also linearized and electroporated into BV-2 cells to generate a stably expressing cell line which was used for experiments.

**Table 1.**
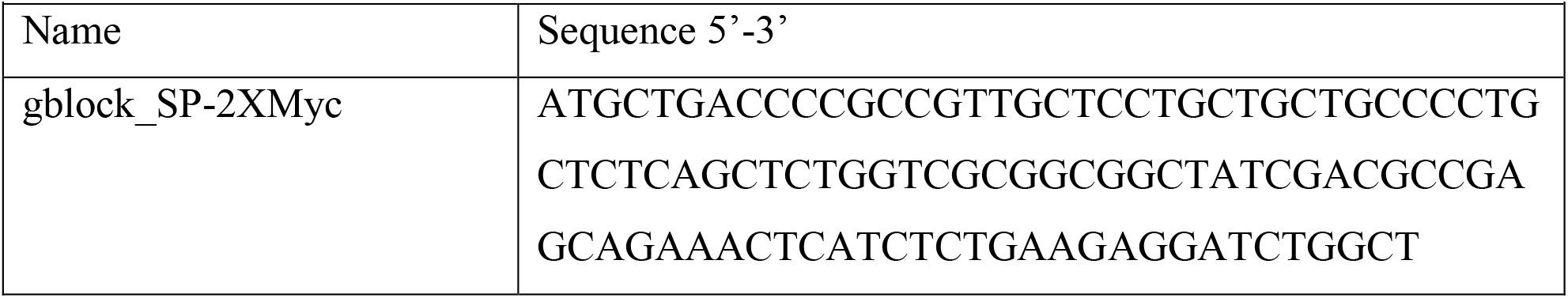
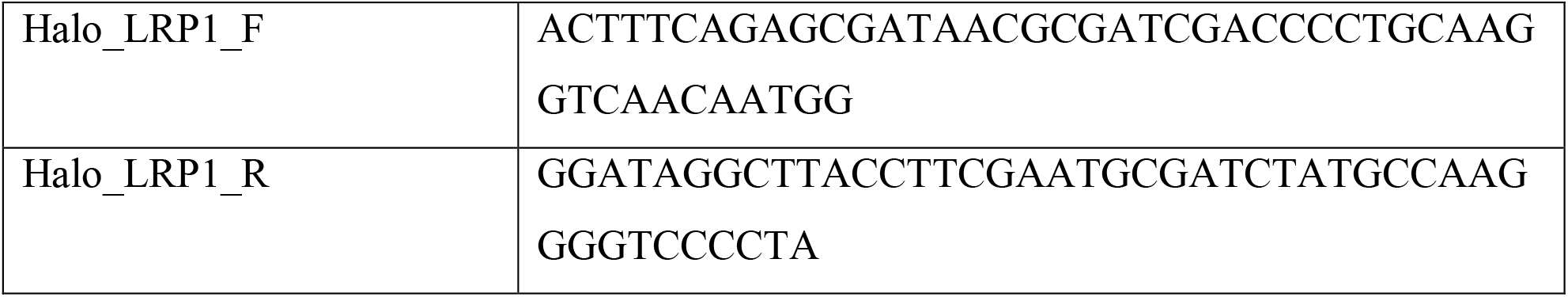
DNA sequences of primers and gblocks used for cloning Halo-mLRP1

### Cell culture

The mouse BV-2 microglial cell line (Goetz Lab, University of Queensland) was sourced from ATCC. BV-2 cells were cultured in RPMI 1640 medium (Thermo Fisher Scientific) supplemented with 10% heat-inactivated fetal calf serum (FCS) (Cytiva) and 2mM L-glutamine (Thermo Fisher Scientific) at 37°C in humidified 5% CO_2_. Cells were seeded at a concentration of 0.1 ×10^6^ cells/mL onto tissue culture treated plates. Transfection was performed using Lipofectamine 2000 (Invitrogen) as per the manufacturer’s instruction.

### Primary cells

Primary co-cultures of microglia and astrocytes were obtained from 2-to 3-day-old CX3CR1^creERT2^ x tdTomato mice; in these mice tdTomato is selectively induced in CX3CR1-expressing cells (i.e. microglia) upon activation of the Cre-ERT2 complex, which subsequently removes a floxed stop codon to initiate expression of the fluorescent reporter (Parkhurst, Yang et al., 2013, Willis et al., 2020). Cells were maintained in DMEM:F12 medium (Thermo Fisher Scientific), supplemented with 100U/mL Pen/Strep (Life Technologies), 2mM Glutamax (Thermo Fisher Scientific), 0.45% glucose (Sigma Aldrich) and 10% FCS. The mixed-glial plates were maintained in T25 tissue culture treated flasks and media was changed every 5 days. To harvest and seed plates for use, the co-culture was washed twice in PBS, gently trypsinized with 0.25% trypsin-EDTA (Invitrogen) for 10 mins in 37°C. Full media was added to deactivate the trypsin and cells were collected into 15mL tubes and spun down at 500g for 5 mins at room temperature. Cells were resuspended in full DMEM:F12 and seeded into MatTek glass bottom dishes (MatTek Corporation) or tissue culture treated plates for experiments.

### Dextran uptake in fixed cells and quantification

BV-2 or primary microglia plated on 12mm glass coverslips were pre-incubated with or without TLR ligands and the coverslips were transferred to a 40μL droplet of fluorescent dextran (OregonGreen,70K MW) made up with or without TLR ligand, to a final concentration of 50μg/mL in RPMI and incubated at 37°C for 15 mins to allow macropinocytic uptake. Cells were then washed in 4°C PBS before fixation for 1 hour with 4% paraformaldehyde. Coverslips were immediately stained with 647-WGA and mounted onto glass slides for viewing. Full Z-stack images were captured on an upright Zeiss AxioImager with Apotome2 and Axiocam 506 camera. Dextran was captured using the Apotome2 whereas 647-WGA was captured using widefield. These images were reformatted as a .tif file with dextran and WGA in the green and red channels, respectively. Reformatting the images is a prerequisite to using automated analysis software. From these images, dextran fluorescence, number of macropinosomes and macropinosome size were measured using a script in ImageJ written and developed by Dr N. Condon (IMB, UQ). This assay was performed three independent times for statistical analysis, where technical replicate data was pooled across the experiments.

For primary microglia, the script was not necessary as images were captured using the Zeiss LSM 880 confocal microscope and not reformatted. Dextran uptake was measured as the Corrected Total Cell Fluorescence (CTCF). CTCF measured the integrated density of dextran fluorescence which is subtracted by the mean background fluorescence of dextran per cell area, as described below. CTCF is thus a standardized measurement of the dextran fluorescence per cell.

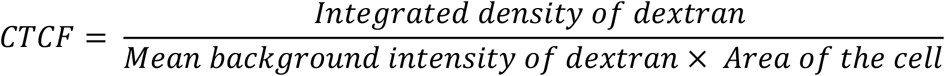

### Live cell microscopy, dextran and marker uptake

Live cell microscopy was performed using an inverted Zeiss LSM 880 confocal with fast airyscan. Cells were seeded at 0.15 × 10^6^ cells/mL in 2 mLs onto MatTek 35mm glass-bottom dishes and imaged in full-RPMI medium in a 37°C heated inset with a cover at 5% CO_2_. To image live dextran uptake, 488-dextran 70K MW (OregonGreen) was prepared to a 2X stock concentration at 100μg/mL. MatTeks were set up to image on the confocal stage and once a field of view was selected, 1.5mL of media was replaced with 500μL of the dextran solution. Imaging commenced immediately to capture initial internalisation of dextran. Live dextran uptake was also paired with live cell imaging of 647-CpG-DNA (300nM) (Integrated DNA Technologies, Singapore) and 488-DQ-BSA (2μg/mL) (Life Technologies).

### Immunoprecipitation

Immunoprecipitation was performed as described previously (Luo et al., 2018). Cells were seeded one day prior to use at 0.2 × 10^6^ cells/mL to a total volume of 10 mL in tissue-culture treated 10cm round dishes (100mm X 20mm TC-treated dish, Corning Costar). The monolayer was washed and collected in 5 mL of PBS and spun down at 500 ×*g* for 5 mins to concentrate the cells in a 15 mL tube for lysis. Lysis buffer (20mM Tris (pH 7.4), 150mM NaCl, 1% NP-40 (sigma), 5mM MgCl_2_, EDTA-free cOmplete protease inhibitors (Sigma Aldrich) and PhosSTOP tablet (Sigma Aldrich)) was used to lyse the cells and lysates were passed through a 27-G syringe to further homogenize the samples. Lysates were centrifuged at 14,000 ×g for 15 mins and the supernatant was collected and used as input. Supernatants were incubated with 2μL anti-LRP1 antibody (Abcam, ab92544) mixed with 20μL IgG sepharose beads for 1 hour at 4°C with rolling for immunoprecipitation. Beads were then washed with lysis buffer 3 times and bound proteins were solubilized in SDS-PAGE sample buffer and boiled at 100°C for 5 mins. Columns were spun at 6, 000 ×g to elute the final sample into 1.5 mL LoBind tubes (525-0130, Bio-Strategy Laboratory Products). MicroSpin columns (27-3565-01, GE Healthcare) were used for all immunoprecipitation and pull downs. Samples were then analyzed by immunoblot.

### Rab8 Activation Assay

Rab Activation Assay was performed as previously described (Tong et al., 2019). Cells were grown the day before in tissue-culture treated 10cm round dishes (100mm X 20mm TC-treated dish, Corning Costar) at a density of 0.2 ×10^6^ cells/mL for a total of 10 mL. After activation with LPS, Poly I:C or CpG-DNA at different time points, the cells were washed and harvested in PBS, spun down at 500 ×*g* for 5 mins and lysed in lysis buffer (25mM Tris pH 7.4, 150mM NaCl, 10mM MgCl_2_, 1% NP-405, 5% glycerol, EDTA-free cOmplete protease inhibitor cocktail and PhosSTOP tablet (Roche Applied Science)) with a 27-G needle. GST-fusion proteins of the Ras-binding domain (RBD) of OCRL (amino acids 539-901) were expressed as recombinant proteins, and they were affinity purified and immobilized on Sepharose beads for pull-down. 400μg/mL protein was loaded into MicroSpin columns (GE Healthcare) with 50μL GST-OCRL-RBD beads and rolled at 4°C for 30 mins. Columns were then washed 3 times with 500μL of lysis buffer and spun down at 600 ×g for 1 min. Samples were eluted by boiling in 2X SDS-PAGE sample buffer for 5 mins and collected in a 1.5mL LoBind tubes by spinning at 6000 ×g for 2 mins. Eluates were then analyzed by immunoblotting.

### Immunoblotting

Cells were lysed in buffer (20mM HEPES (pH7.4), 100mM NaCl, 1% NP-40 (Sigma Aldrich) with added cOmplete protease inhibitor cocktails (Sigma Aldrich) and phospho-STOP tablets (Sigma Aldrich) and lysates standardized with SDS-PAGE sample buffer using the Pierce BCA Protein Assay Kits as per the manufacturer’s instructions. Samples were run on 10% acrylamide gels in running buffer (0.25M Tris, 1.92M Glycine and 1% SDS). Proteins were transferred to a polyvinylidene difluoride membrane (Biotrace) which was incubated for 1 hour with 5% skim milk in TBS-Tween (TBST) buffer to block non-specific protein binding. Membranes were then incubated with primary antibodies at 4°C overnight with shaking. After 3 washes, 10 mins each with TBST, membranes were incubated with secondary antibodies at room temperature for 1 hour. Membranes were washed again and developed with SuperSignal™ WestPico chemiluminescent reagent (Thermo Fisher Scientific) according to the manufacturer’s instructions. Blots were imaged on the Bio-Rad Chemi-Doc Imaging System.

### Quantitative reverse transcription PCR

BV-2 were seeded onto 12-well tissue-culture treated plates (sterile, 12 well size TC-treated plate, Corning Costar) at a density of 0.2 ×10^6^ cells/mL (1 mL per well) in complete medium overnight. Cells were incubated with DMSO or IPI-549 (100nM) for 1 hour prior to activation with Poly I:C (10μg/mL) or CpG-DNA (300nM). RNA was collected at 2-, 6- and 24-hours post activation as well as unactivated cells as a baseline control. RNA was extracted using Trizol™ (Life Technologies) as per manufacturer’s instruction. RNA was resuspended in 20μL in DNAse, RNAse free dH_2_O. RNA concentration was quantified using NanoDrop and samples were standardized to the same concentration. cDNA was synthesized from the RNA template using SuperScript™ III First-Strand Synthesis System for RT-PCR (Life Technologies) as per manufacturer’s instruction. cDNA was diluted to a final volume of 200μL in dH_2_O. qPCR was performed using SYBR Green (Applied Biosystems) with optimized qPCR primers. HPRT (Hypoxanthine-guanine phosphoribosyltransferase) was used as a housekeeping control. Delta CT and fold change (2^-deltaCT^) was calculated using Microsoft Excel and was analyzed using GraphPad Prism 8.

### AlphaLISA immunoassay for the detection of IL-6 and IL-10

Cells were pre-incubated for 1 hour with DMSO (untreated control) or IPI-549 (100nM) to inhibit PI3Kγ. Cells were then activated with CpG-DNA (300nM) and the conditioned media was collected at 6 hours and 24 hours post activation. Media samples were kept on ice and used immediately for AlphaLISA in a 384-well plate as per manufacturer’s instruction. The plate was read with excitation and emission at 680nm and 615nm, respectively Data was analyzed using GraphPad Prism 8.

### Statistical Analysis

Statistical analysis was performed using GraphPad Prism software package version 8. Data was presented as mean ± S. D. Data used for statistical analysis was collected and pooled from a minimum of n=3 independent experiments, unless otherwise stated. Data sets were assessed for significance by two-tailed Student’s T-test, Mann-Whitney U test (for non-normal distributions), ordinary one-way or two-way ANOVA (with additional posthoc testing, described in figure legends). Significance was considered if p<0.05 (* p<0.05, ** p<0.01, *** p<0.001, **** p<0.0001).

## Acknowledgements

The authors thank Tatiana Khromykh (UQ IMB) and Dr Emily Willis (UQ QBI) for technical assistance with molecular cloning and primary microglia respectively. Janelia Fluor Halo probes were generously provided by Luke Lavis (HHMI Janelia). The TagRFP-Akt construct was kindly provided by Dr James Burchfield (USyd). The LRP1 minireceptor used for cloning was provided by Prof Maria-Paz Marzolo (Pontifical Catholic University of Chile). Imaging was performed in IMB’s Microscopy Facility incorporating the Cancer Ultrastructure and Function Facility funded by Australian Cancer Research Foundation. YH and NT were supported by PhD scholarships from the Australian government and YH was awarded funding from the Yulgilbar Alzheimer Research Program. Z-JX was supported by a Chinese Academy of Science Scholarship. SJT was supported by a UQ/IMB Postgraduate Scholarship. JV holds a Senior Medical Research Fellowship from the Sylvia and Charles Viertel Foundation. Salary and research funding was from the National Health and Medical Research Council of Australia (JLS APP1176209; APP108511), the National Health and Medical Research Council (NHMRC; Project Grant 1124503 and NHMRC; Project Grant 1124503) and the Australian Research Council (LL DE180100524; JLS, AAW DP180101910; JV DE150101578).

## Author contributions

YH, AAW and JLS conceptualized the project and designed experiments. YH and JLS wrote the manuscript. YH, NT, Z-JX and SJT performed experiments and contributed data. LL and JV provided reagents, cells and expert advice.

## Data availability section

This study includes no data deposited in external repositories.

## Appendix Figure Legends

**Appendix Figure S1.**
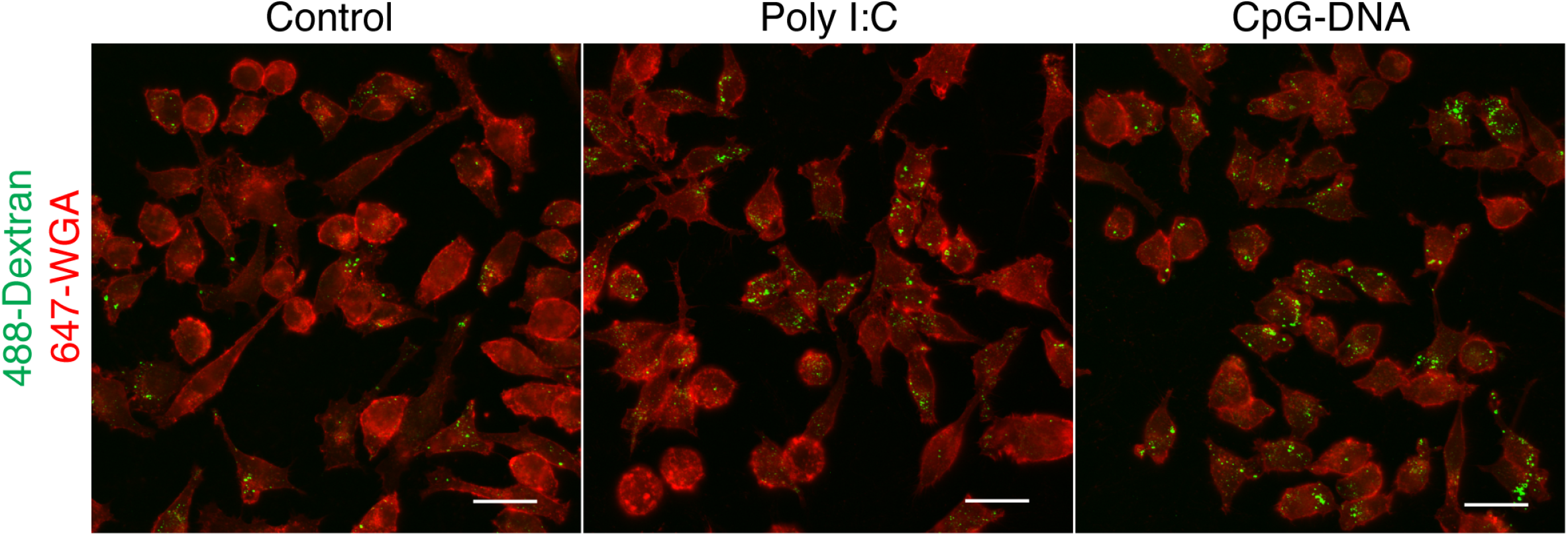
Endosomal TLR activation upregulates macropinocytosis. Representative fixed cell staining of BV-2 microglia activated with Poly I:C (10μg/ml) or CpG (300nM) for 1 hour and incubated for 30 mins with 488-Dextran 70K MW (green). Control cells were unstimulated prior to the 30 mins incubation with dextran. Images are max intensity projections. All scale bars represent 20μm.

**Appendix Figure S2.**
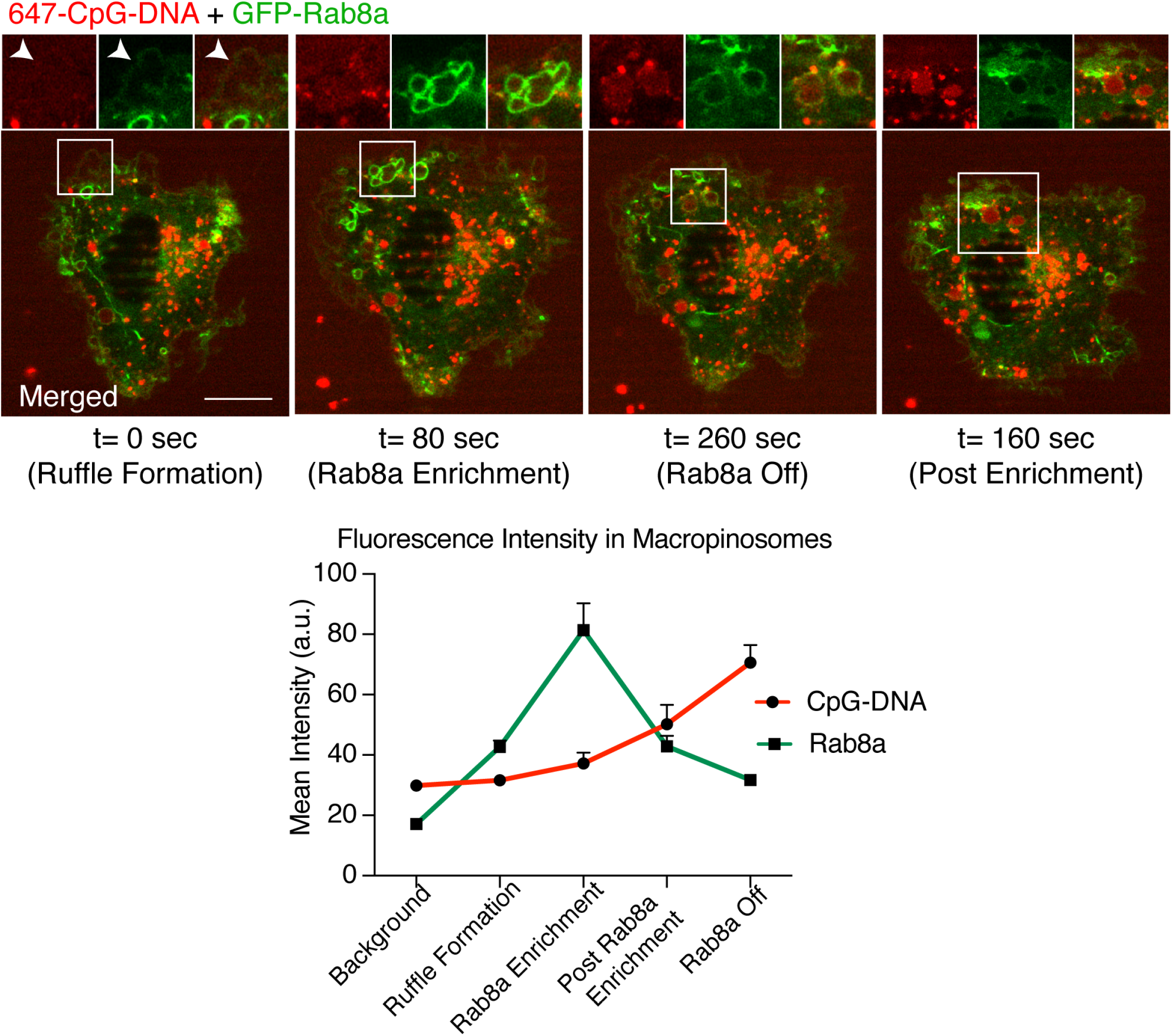
Rab8a is enriched on macropinosomes which internalize TLR9 ligand, CpG-DNA. Live cell confocal imaging of BV-2 microglia transiently transfected with GFP-Rab8a (green). Cells were incubated with 647-CpG-DNA (red) and images were captured over time. Movie sequence depicts macropinosome formation, 647-CpG-DNA accumulation and GFP-Rab8a enrichment. Fluorescence intensity of 647-CpG-DNA and GFP-Rab8a were measured within macropinosomes and plotted on graph. Measurements were pooled from technical replicates with n=3 independent experiments. Scale bar represents 10μm.

**Appendix Figure S3.**
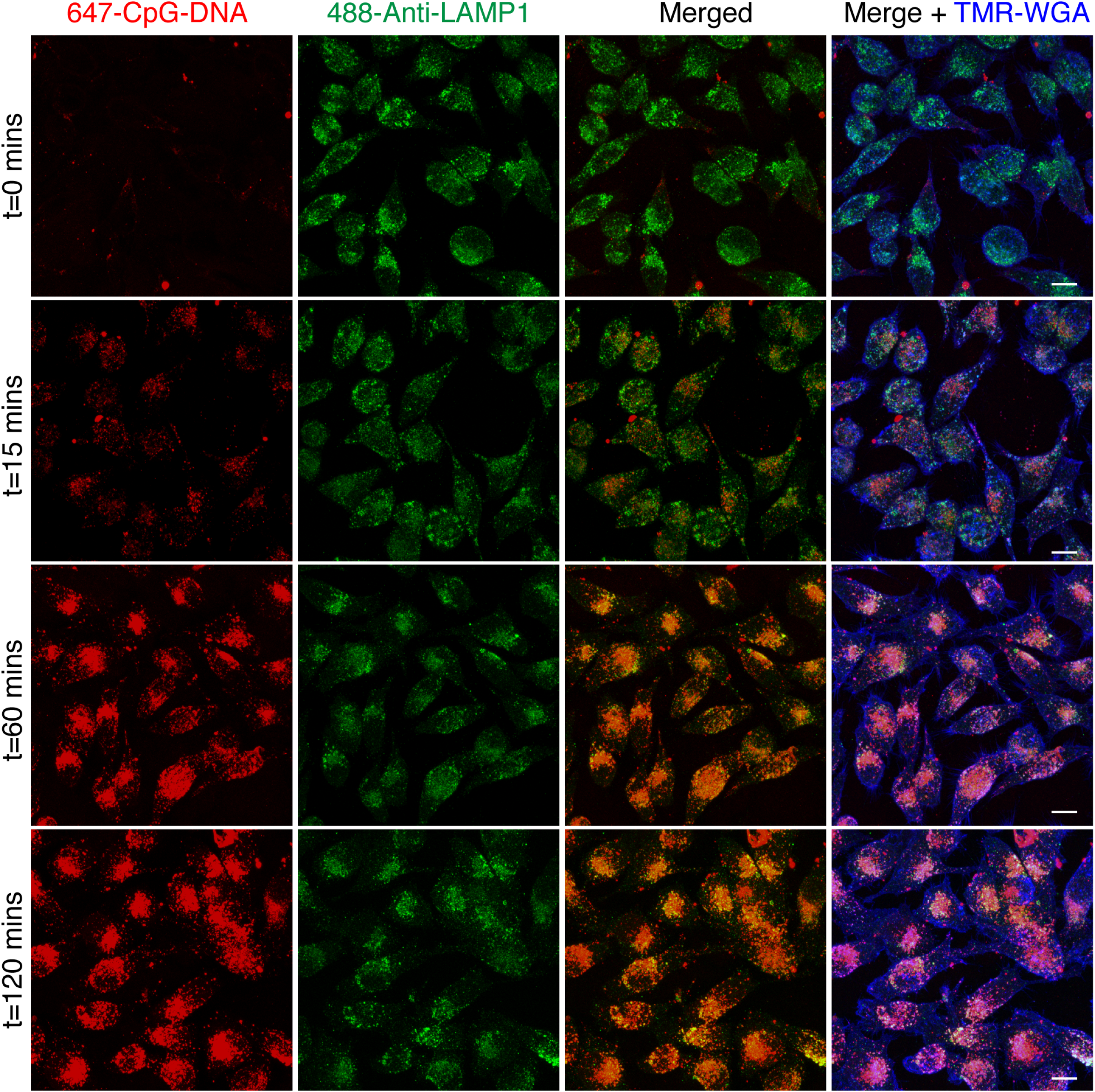
TLR9 ligand, CpG-DNA is trafficked to LAMP1 positive lysosome at later time points. Representative staining of BV-2 cells incubated with 647-CpG-DNA (red) and fixed across a 2 hour time course. Cells were then stained with anti-LAMP1 antibody (green) and wheat germ agglutinin was used to label the cell outline (blue). As 647-CpG-DNA accumulates in the cell, it co-localizes strongly with anti-LAMP1 staining. Scale bar represents 10μm.

**Appendix Figure S4.**
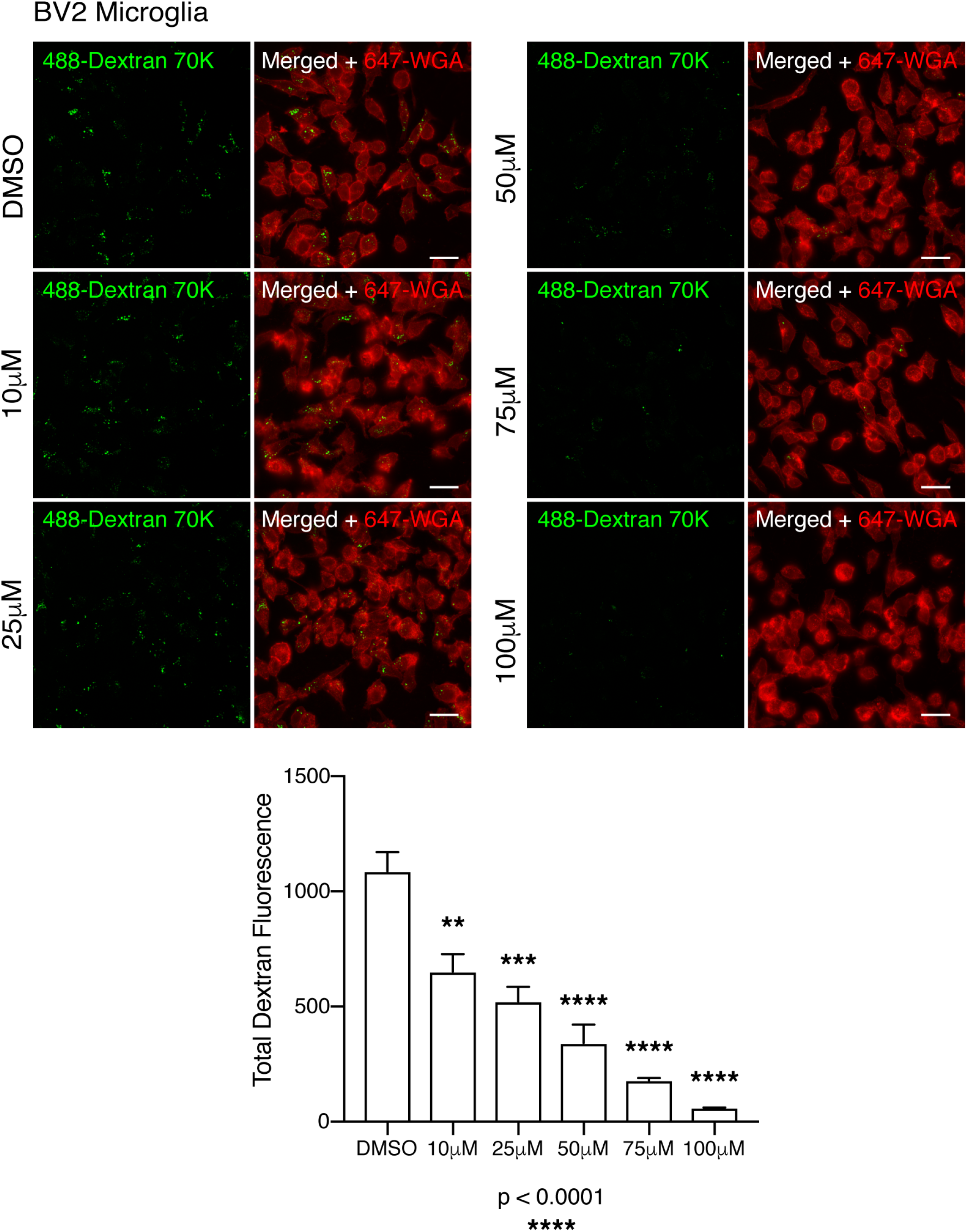
EIPA inhibition of macropinocytosis and dextran uptake. BV-2 microglia pre-treated for 1 hour with increasing concentrations of EIPA (10-100μM) and DMSO as a control. Cells were then incubated with 488-Dextran 70K MW (green) for 30 mins to allow for uptake, then fixed and stained with wheat germ agglutinin (WGA) red) to label the cell outline. Total dextran fluorescence was measured by hand and displayed as mean ± SD. n=3 independent experiments, \*\*\*\**p<0*.*0001* for interaction (ordinary one-way ANOVA with Dunnett’s multiple comparison’s test, significance between DMSO control and drug treatment displayed on graph). All scale bars represent 20μm.

**Appendix Figure S5.**
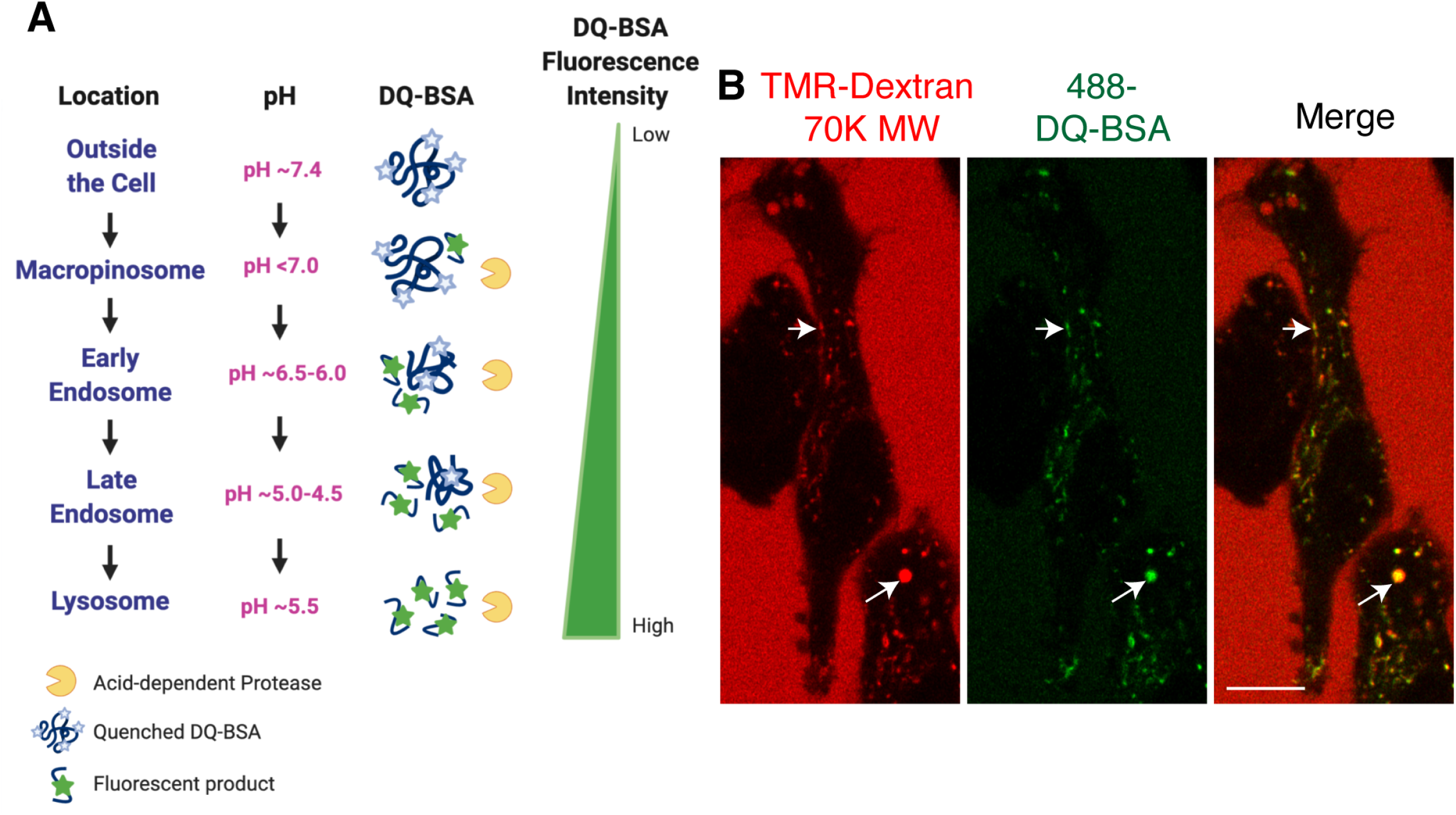
DQ-BSA fluorescence in acidic endosomes in microglia. **A**. Diagram demonstrating DQ-BSA fluorescence in endosomes. DQ-BSA is quenched under physiological pH and its breakdown by acid-dependent proteases produces a fluorescent product. Single time point of live cell confocal imaging of BV-2 microglia incubated with TMR-Dextran 70K MW (red) and DQ-BSA (green). DQ-BSA fluoresces under proteolytic cleavage in acidic endosomes, co-labelled by the fluid-phase marker, TMR-dextran (arrows). Scale bar represents 10μm.

**Appendix Figure S6.**
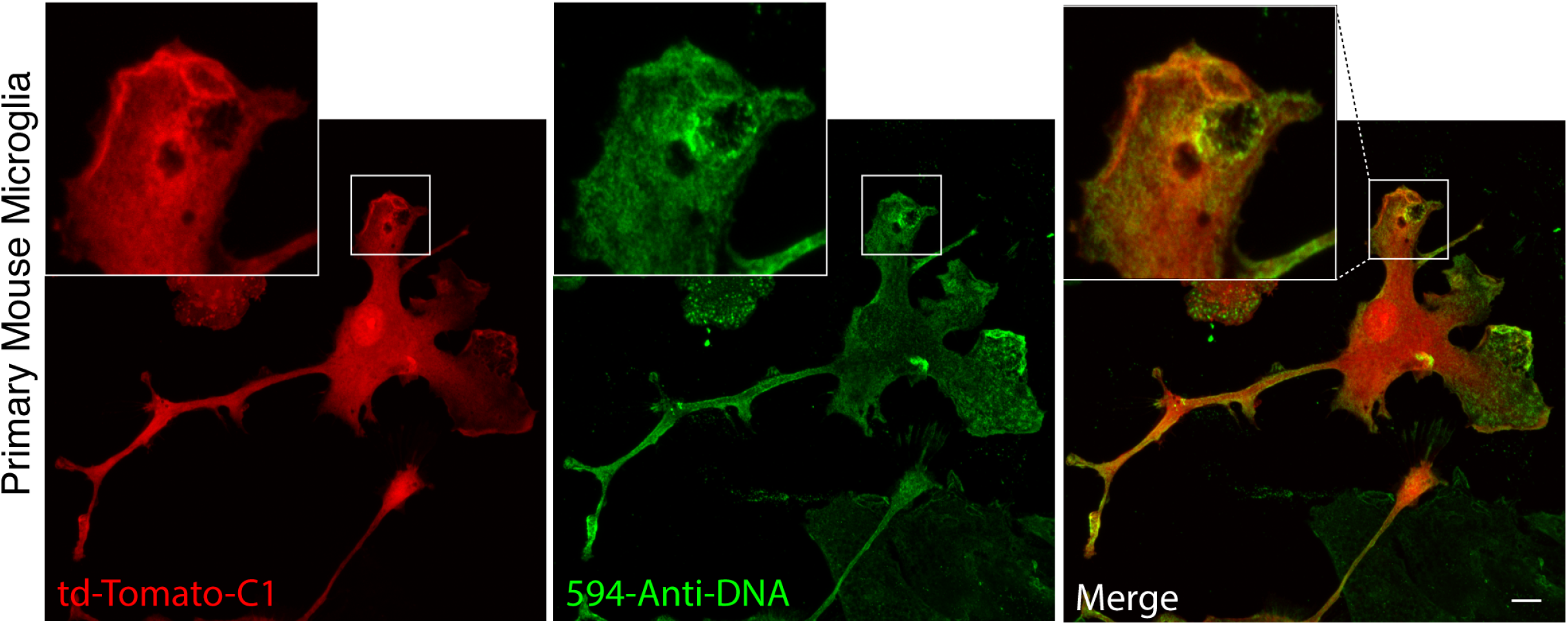
DNA debris is localized around macropinosomes. Representative fixed confocal imaging of primary microglia selectively expressing tdTomato-C1 (green). Cells were left unpermeabilized and stained with anti-DNA antibody to mark exogenous DNA (green). Scale bar represents 10μm.

**Appendix Figure S7.**
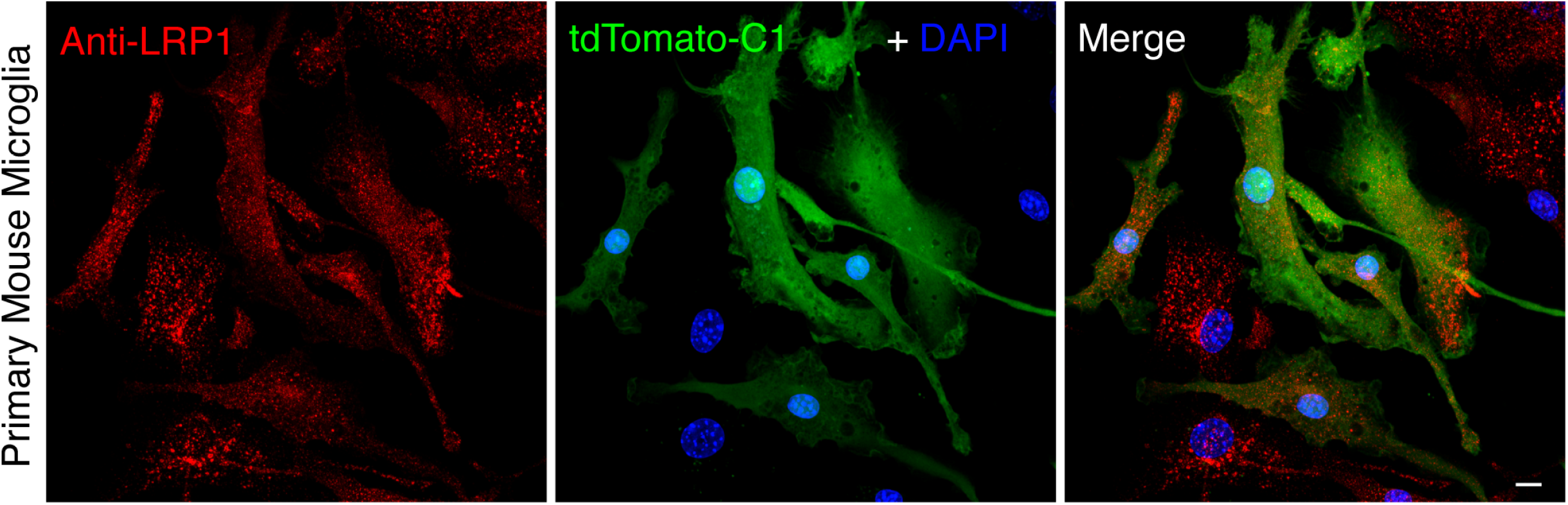
LRP1 is localized to recycling endosomes in microglia. Representative confocal imaging of fixed primary microglia selectively expressing tdTomato-C1 (green). Cells were stained with anti-LRP1 antibody to mark endogenous LRP1 (red). Microglia in astroglial co-cultures, all cell nuclei are labelled with DAPI. LRP1 staining in the astrocytes can be seen by DAPI labelling and the absence of tdTomato-C1. Scale bar represents 10μm.

**Appendix Figure S8.**
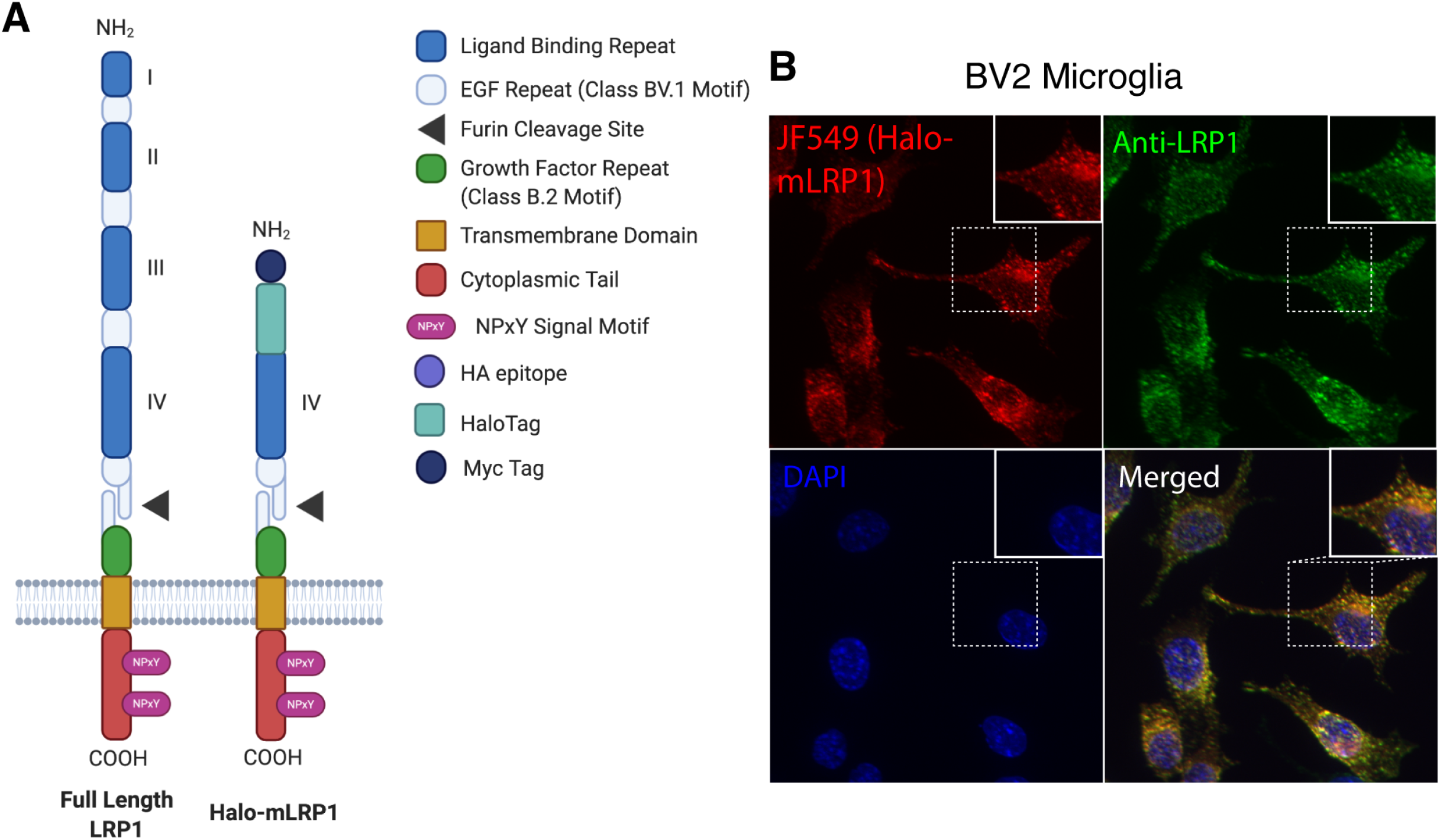
Halo-mLRP1 design and expression. **A**. Diagram of full length LRP1 (600kDa) in comparison to the Halo-tagged minireceptor (222kDa), used for live cell imaging. Halo-mLRP1 contains only the 4^th^ ligand binding domain and the exofacial HaloTag. Diagram not to scale. **B**. Fixed cell imaging of BV-2 microglia stably expressing Halo-mLRP1. Cell permeable halo-ligand (JF549) marks Halo-mLRP1 and the anti-LRP1 labels all LRP1 including Halo-mLRP1 and endogenous LRP1. DAPI (blue) labels cell nucleus. Scale bar represents 10μm.

